# IRAK1-mediated coincidence detection of microbial signals licenses inflammasome activation

**DOI:** 10.1101/2019.12.26.888776

**Authors:** Sharat J. Vayttaden, Margery Smelkinson, Orna Ernst, Rebecca J. Carlson, Jing Sun, Clinton Bradfield, Michael G. Dorrington, Jonathan Liang, Nicolas Bouladoux, Rachel A. Gottschalk, Kyu-Seon Oh, Gianluca Pegoraro, Sundar Ganesan, Dominic De Nardo, Eicke Latz, Yasmine Belkaid, Rajat R. Varma, Iain D.C. Fraser

## Abstract

The innate immune system signals through various higher order signaling complexes called supramolecular organizing centers (SMOCs), which typically organize components of a single pathway. While innate immune signaling pathways have been largely characterized using single receptor stimuli, responses to pathogens require the coordinated engagement of multiple pathways. Here, we report an IRAK1-containing SMOC formed specifically when multiple receptors are activated, which recruits select components of the TLR, MAPK and inflammasome pathways. This allows for signal flux redistribution from TLRs to inflammasomes and facilitates inflammasome licensing through an MKK7-JNK axis, which is defective in *Irak1*^−/−^ mice. Furthermore, this defect in *Irak1*^−/−^ mice manifests in increased susceptibility to inflammasome-sensitive pathogens and diminished IL1 production from inflammasomes after co-TLR priming. Thus, IRAK1 SMOCs form a multi-pathway coordinating hub for coincidence detection of microbial signals, which may be employed by innate immune cells as a threat assessment and thresholding mechanism for inflammasome activation.

## Introduction

Pattern recognition receptor (PRR) families are expressed on sentinel cells of the innate immune system like macrophages. These receptor families detect conserved pathogen-associated molecular patterns (PAMPs) on microbes, and also intrinsic damage-associated molecular patterns (DAMPs) encountered during tissue damage (Akira et al., 2006; Janeway and Medzhitov, 2002; Mishra et al., 2017). Upon activation by their cognate ligands, these receptors promote inflammatory gene expression or cell death that can facilitate infection clearance and restoration of tissue homeostasis. The molecular details of PRR signaling pathways have been predominantly derived from studies that involve stimulation of single PRRs by their cognate ligands (Alexopoulou et al., 2001; Brown et al., 2002; Fitzgerald et al., 2001; Gottar et al., 2002; Hemmi et al., 2000; Horng et al., 2002; Jurk et al., 2002; McCoy and O’Neill, 2008; Shimazu et al., 1999; Takeuchi et al., 2001; Takeuchi et al., 2002; Watson et al., 1977; Wright et al., 1990; Wright et al., 1989). While these studies have revealed that there are common frameworks for organization of the PRR pathways, few have addressed how simultaneously activated PRR pathways interact to ensure an appropriate host response to the combined PAMP and/or DAMP signals encountered during microbial infection or tissue damage (Bagchi et al., 2007; Gottschalk et al., 2019; Lin et al., 2017; Liu et al., 2015; Napolitani et al., 2005; Suet Ting Tan et al., 2013; Tan et al., 2014).

Toll-like Receptors (TLRs) are an extensively studied class of PRRs. Once activated, all TLRs can signal through IRAK and/or TRAF proteins, eventually leading to activation of transcription factors from the AP1, NF-κB and IRF families, which regulate the gene transcriptional programs initiating an immune response (O’Neill, 2006). Since in a physiological setting immune cells encounter a cocktail of TLR ligands, there are likely mechanisms to encode a multi-TLR response that are distinct from responses observed from activation with single TLR ligands. Additionally, the host may seek to avoid a full-fledged immune response to very low levels of single TLR ligands that pose no pathogenic threat, while the presence of multiple PRR ligands may exceed a ‘threat assessment’ threshold and warrant a stronger response (Evavold and Kagan, 2019). This would require some form of coincidence detection to function within the PRR pathways. Given the commonalities in downstream signaling from different TLRs, signal integration strategies must also account for use of shared signaling components, but it remains unclear how this process might be regulated to achieve appropriate levels of innate immune activation (Franz and Kagan, 2017; Kawai and Akira, 2011).

PRR signaling can be organized through formation of macromolecular protein complexes (Bryant et al., 2015), and higher-order signaling complexes have been termed supramolecular organizing centers (SMOCs) (Kagan et al., 2014; Qiao and Wu, 2015) or signaling through cooperative assembly formation (SCAF) (Vajjhala et al., 2017). These complexes facilitate increased local concentrations of signaling components to promote weak interactions. They often consist of sensors that detect the activation status of PRRs and promote context-specific downstream effector responses. The FAS-DISC (Algeciras-Schimnich et al., 2002), PIDDosomes (Park et al., 2007), myddosomes (Lin et al., 2010; Motshwene et al., 2009), putative trifosomes (Bryant et al., 2015), RLR complexes (Hou et al., 2011) and inflammasomes (Lu et al., 2014) are examples of SMOCs or SCAFs associated with PRR signaling. Most known SMOCs of the innate immune response are associated with a single pathway and haven’t been shown to coordinate signaling of multiple pathways. Since the SMOCs studied so far are elicited by single PRR activation, it is possible that multi-PRR stimulation induces additional SMOCs that are assembled in response to combined input signals, and that the resulting crosstalk facilitates a more effective immune response.

TLR2 and TLR4 are well-studied PRRs known to play cooperative roles in innate immune responses (Jeon et al., 2017; Zhang et al., 1997). These TLRs, when individually stimulated, signal through either MyD88 alone or a combination of the MyD88 and TRIF signaling pathways. Myddosomes formed under single TLR stimulation in mouse macrophages typically include a complex of MyD88, IRAK4 (Motshwene et al., 2009) and IRAK2 (Lin et al., 2010). It has been suggested that IRAK1 and IRAK2 are both involved in initial TLR signaling, while IRAK2 plays a critical role in sustaining the TLR response (Kawagoe et al., 2008). In murine macrophages, single ligand-induced TLR signaling and cytokine responses are strongly dependent on IRAK2, while IRAK1 has a more limited role (Sun et al., 2016). Since these studies employed single TLR ligands, it remains unclear whether signaling mechanisms induced through single-TLR stimulation are comparable to those induced by multi-TLR stimulation.

Here, we describe an IRAK1-containing SMOC that is preferentially induced by combined activation of TLR4 and TLR1/2 by either PAMPs or DAMPs or through multi-PRR activation by bacteria. These IRAK1 SMOCs lacked myddosome components but were enriched for certain components of the TLR, MAPK and inflammasome pathways, namely pTBK1, TRAF6, pMKK7, pJNK, ASC, NLRP3 and NLRC4. Inflammasome responses as measured by IL-1α and IL-1β secretion following co-TLR priming were perturbed in *Irak1^−/−^* macrophages. However, co-TLR activated transcriptional priming of murine inflammasome genes was unaffected by IRAK1 deficiency. *Irak1^−/−^* mice showed increased susceptibility to *Yersinia pseudotuberculosis* infection and dysregulated serum cytokine levels. *Irak1^−/−^* mice with respiratory *Pseudomonas aeruginosa* infection showed decreased IL-1α and IL-1β in the bronchoalveolar lavages. These data demonstrate a previously unappreciated role for IRAK1 in forming a SMOC induced specifically through coincident detection of multiple PAMPs which facilitates licensing of the inflammasome response in a JNK-dependent process.

## Results

### IRAK1 forms discrete intracellular clusters in response to multi-PRR activation

*Irak1^−/−^* mice have been shown to be only slightly protected in single TLR stimulated endotoxic shock (Swantek et al., 2000) at a dose of LPS where 60% WT mice survive. This is in contrast to knockouts of other early TLR signaling components like MyD88, TRIF, IRAK4, and IRAK2 (Kawai et al., 1999; Suzuki et al., 2002; Tang et al., 2018; Wan et al., 2009), where the knockout mice show strong protection from endotoxemia at LPS doses that are lethal to all but 0-10 % of WT mice. We subjected *Irak1^−/−^* mice to endotoxic shock at an LPS dose that achieved 90% mortality in WT mice and the survival of WT and *Irak1^−/−^* mice were not significantly different (Fig.1A). This suggested that IRAK1 might be largely dispensable for LPS-induced responses. We then tested if the gene transcription response for a panel of TLR-induced genes was different between WT and *Irak1^−/−^* murine bone marrow-derived macrophages (BMDMs) for three different TLR ligands, namely P3C (TLR1/2), PIC (TLR3) and Kdo-2 Lipid A (TLR4). These ligands were selected for their known ability to signal through either MyD88 (P3C) or TRIF (PIC) or both (Kdo-2 Lipid A). The gene transcription profile showed high similarity between WT and *Irak1^−/−^* mice (Fig. 1B, Table S1). This agrees with the prior observation that, in contrast to other proximal TLR signaling components, IRAK1 knockout has minimal effect on single TLR ligand-induced cytokine responses in murine BMDMs (Sun et al., 2016). This contrasts TLR7- and TLR9-mediated IFNα induction in murine plasmacytoid dendritic cells, which is strongly IRAK1-dependent (Uematsu et al., 2005).

**Figure 1.**
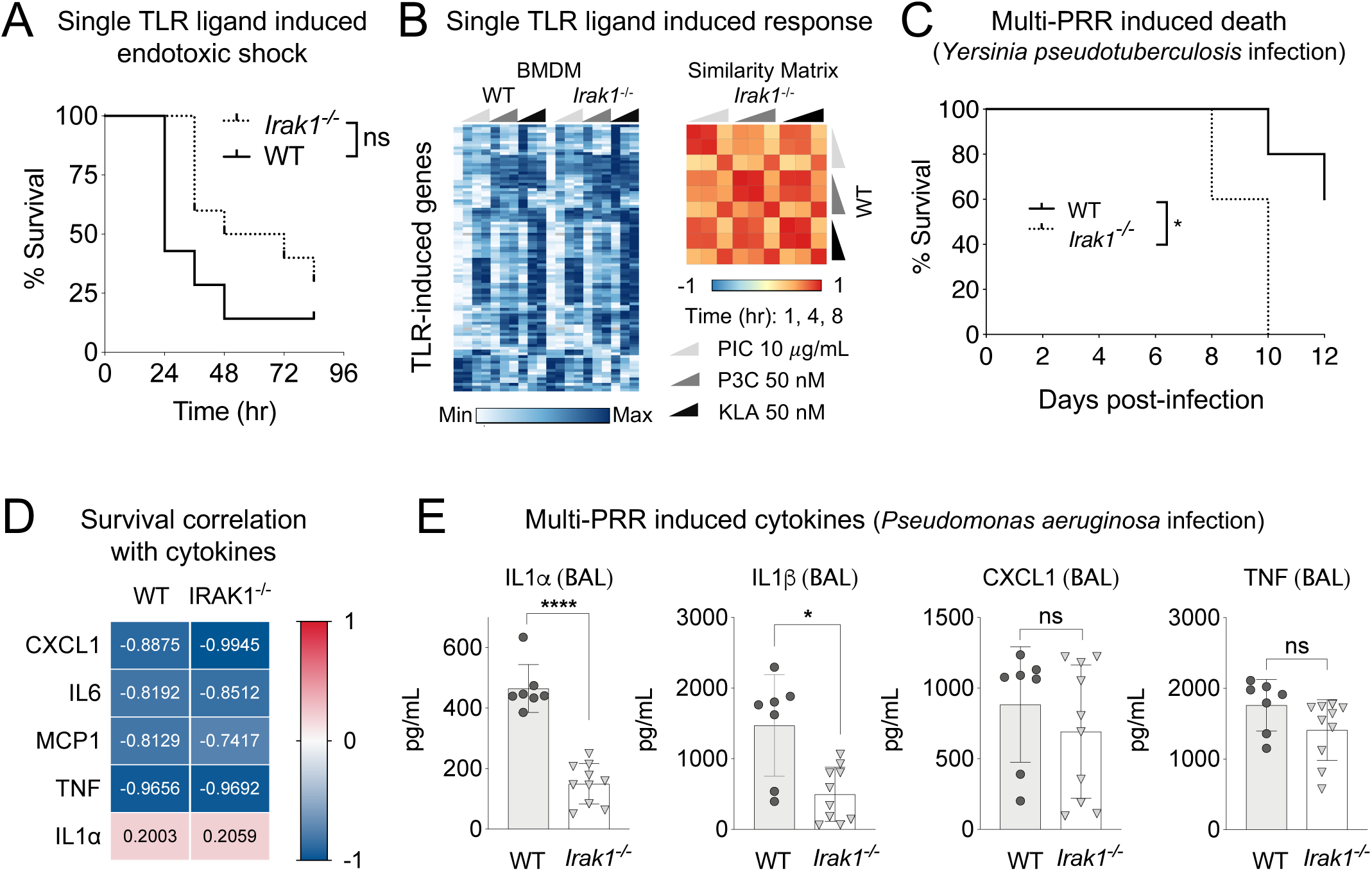
IRAK1 has different roles in single vs multi-TLR signaling. (A) Survival after intraperitoneal injection of LPS in WT and *Irak1*^−/−^ mice. Log-Rank Mantel-Cox Test. (B) Transcriptional response to single TLR ligands Poly(I:C), P3C and Kdo-2 Lipid A in BMDMs 0, 1, 4 and 8 h after stimulation. Median from two experiments. Similarity Matrix – Pearson’s Correlation Coefficient. mRNA levels were assayed by Fluidigm microfluidic qPCR. See Table S1. (C) Survival after infection with *Yersinia pseudotuberculosis* in WT and *Irak1*^−/−^ littermate mice. Log-Rank Mantel-Cox Test. (D) Survival correlation with Day 3 serum cytokine levels in mice infected with *Yersinia pseudotuberculosis*. Pearson’s Correlation Coefficient (E) Cytokines from broncho-alveolar lavages of WT and *Irak1*^−/−^ mice 16 h after intranasal injection of *Pseudomonas aeruginosa*. (Panel E) Data are represented as mean ± SD. Unpaired t test with Welch’s correction. p = 0.1234 (ns), 0.0332(*); 0.0021 (**); 0.0002 (***); < 0.0001 (****). Data shown are representative of at least two independent experiments.

IRAK1 is ubiquitously expressed in multiple tissues, and phylogenetic analysis suggests that IRAK1 has been conserved since early vertebrates (Gosu et al., 2012). The tissue ubiquity and evolutionary conservation of IRAK1 suggests that IRAK1 serves important functions that might not be evident in certain single TLR stimulation studies. We sought to test if under more complex stimulation, such as a multi-PRR response brought about by bacterial infection, *Irak1^−/−^* mice would fare differently than WT mice. For this we used *Yersinia pseudotuberculosis* which is known to infect macrophages and dendritic cells in mice (Brodsky and Medzhitov, 2008; Fonseca et al., 2015). We found that *Irak1^−/−^* mice showed substantially increased mortality compared to WT mice (Fig. 1C). Serum IL1α levels at day 3 were positively correlated with length of survival in direct contrast to other inflammatory cytokines (Fig. 1D). This is consistent with the critical role for inflammasome-dependent IL-1 responses in host defense against *Yersinia* species, and the targeting of the inflammasome pathway by *Yersinia* T3SS effectors (Chung and Bliska, 2016). Due to the survival correlation of IL1 in the *Yersinia* model, we tested the IRAK1-dependency of cytokine responses in a more acute but localized model of *Pseudomonas aeruginosa* lung infection. Broncho-alveolar lavages showed that *Irak1^−/−^* mice had markedly reduced IL-1α and IL-1β protein levels, but no significant change in CXCL1 or TNF responses (Fig. 1E).

To investigate the effect of a multi-PRR stimulation on intracellular localization of IRAK1, we challenged an *Irak1^−/−^* immortalized bone marrow derived macrophage (iBMDM) cell line stably expressing an IRAK1-mCherry fusion with live *Salmonella enterica* serovar Typhimurium expressing GFP (Fig. 2A, Fig. S1A, Movies 1-2). Multi-PRR stimulation led to a change in the spatiotemporal pattern of IRAK1, from diffuse in untreated cells, to discrete clusters upon challenge with bacteria. This IRAK1 clustering response could be recapitulated with a panel of heat-killed Gram negative and Gram positive bacteria (Fig. 2B) or on co-stimulation with TLR4 and TLR1/2 ligands in iBMDMs (Fig. 2C, D, Fig. S1B-D, Movies 3, 4), but was diminished in cells treated with single TLR4 or TLR1/2 ligands (Fig. 2D, S1C, Movies 5, 6). We confirmed that the IRAK1-mCherry clusters were recognized by IRAK1-specific mouse antibodies (Fig. S1E), and that this staining was absent in *Irak1^−/−^* iBMDMs (Fig. S1F). The pixel location and intensities of mCherry and the Alexa Fluor 488-labeled IRAK1 antibody had a good correlation (Fig. S1G, H). Pearson’s Correlation Coefficient (PCC) was used for describing the correlation of the intensity distributions between channels, and Mander’s Colocalization Coefficient (MCC) for indicating an overlap of the signals thus representing the two major metrics of colocalization (Dunn et al., 2011). Using these criteria, we see that IRAK1 clusters also colocalized strongly with the IRAK1 rabbit antibody staining (Fig. S1I, J).

**Figure 2.**
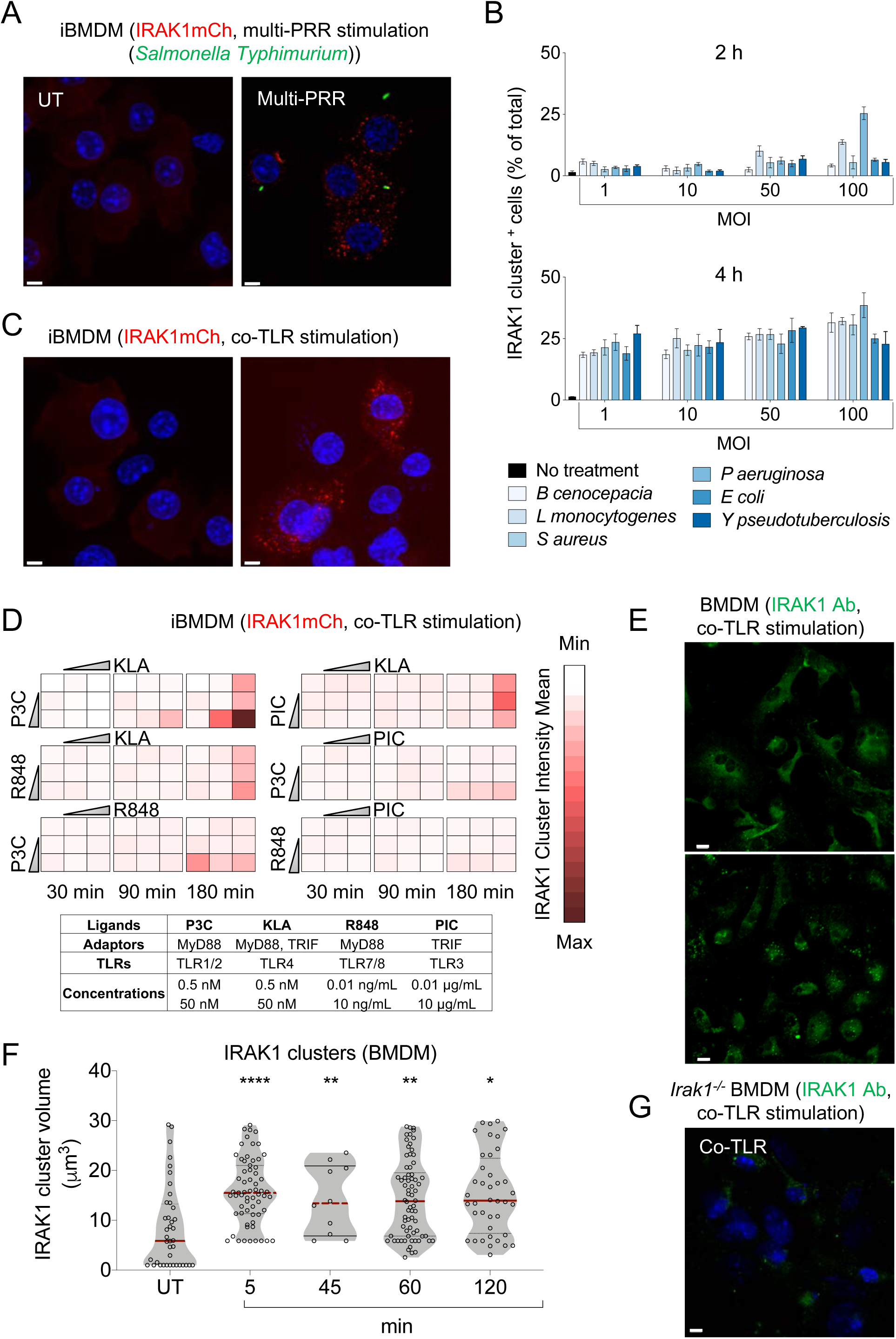
IRAK1 forms discrete intracellular clusters in response to multi-PRR activation. (A) Images of IRAK1 in iBMDMs 0, and 2h after infection with GFP expressing *Salmonella enterica* serovar Typhimurium (MOI10). Scale Bar: 10*μ*. (B) Quantification of IRAK1 clusters in iBMDMs on multi-PRR stimulation resulting from treatment with heat-killed Gram positive and Gram-negative bacteria for 0, 2 and 4h. Data are represented as median ± MAD (C) Images of IRAK1 in iBMDMs 0 and 2 h after co-stimulation with P3C and Kdo-2 Lipid A (50 nM each). Scale Bar: 10*μ*. (D) Quantification of IRAK1 clustering in iBMDMs 0, 30, 90, and 180 min after pairwise stimulation with TLR ligands – Kdo-2 Lipid A, P3C, Poly(I:C) and R848. Data are represented as mean (n = 3). IRAK1 antibody (Proteintech 10478-2-AP) staining in (E, F) BMDMs and (G) *Irak1^−/−^* BMDMs on co-stimulation with P3C and Kdo-2 Lipid A (500 nM each). Scale Bar: 10*μ*. (Panel F) Tamhane’s T2 multiple comparisons test. p = 0.1234 (ns), 0.0332(*); 0.0021 (**); 0.0002 (***); < 0.0001 (****). Data shown are representative of at least two independent experiments. See also supporting Fig. S1, Movies 1-5.

To determine whether IRAK1 clustering is induced by other TLR ligand pairs, we tested a panel of four different TLR ligands across a dose range. IRAK1 clusters were detectable with all tested co-TLR stimulations (Fig. 2D), but were most strongly induced by co-activation of TLR4 and TLR1/2 with the strongest separation between single vs co-TLR stimulated IRAK1 clustering being observable at 90 min. We therefore chose this ligand pair and time point to further investigate co-TLR-induced IRAK1 clusters. This ligand pair also induced clustering of endogenous IRAK1 in primary BMDMs (Fig. 2E, F) and this staining was absent in *Irak1^−/−^* BMDMs (Fig. 2G). Since IRAK1 clustering was strongest on co-stimulation of TLR4 and TLR1/2, we sought to determine if other ligands known to co-stimulate these receptors would induce IRAK1 clustering. Biglycan, a small leucine-rich proteoglycan can be found in two distinct forms, an extracellular matrix (ECM) bound form and a soluble form. The soluble form of biglycan is released from the ECM upon tissue damage and can function as a danger-associated molecular pattern (DAMP) through activation of both TLR4 and TLR2 (Erridge, 2010; Merline et al., 2011; Schaefer et al., 2005). We observed a substantial increase in IRAK1 clustering with the soluble biglycan DAMP, but it peaked later than the KLA and P3C PAMP combination, and the biglycan-induced cluster size appeared smaller (Fig. S1K). Thus, it seems that IRAK1 clustering can be induced with multi-PAMP or complex DAMP stimulation.

### Properties of IRAK1-containing clusters

Given the preferential occurrence of IRAK1 clusters in response to co-stimulation of TLRs, we sought to characterize their properties and potential role in TLR signaling. There have been reports of IRAK1 ubiquitylation and phosphorylation leading to degradation in different cell types and contexts (Gottipati et al., 2008; Schauvliege et al., 2006; Yamin and Miller, 1997). It has also been suggested that IRAK1 antibodies can fail to detect their epitopes on account of such modifications, which can lead to challenges in interpretation of diminished band intensities on western blots (Emmerich and Cohen, 2015). To first investigate whether IRAK1 clusters were associated with protein degradation machinery, we co-stained for the 26S proteasome, but observed no substantial colocalization (Fig. 3A). The SCF (Skp1-Cullin1-F-box)-β-TrCP complex has been identified as the E3 ligase responsible for K48-linked ubiquitination of IRAK1 (Cui et al., 2012), while the Pellino proteins have been shown to mediate K63-linked polyubiquitination of IRAK1 after IL-1 stimulation (Schauvliege et al., 2006). We tested to see if the IRAK1 clusters were colocalized with β-TrCP (Fig. S2A) or Pellino (Fig. S2B) under co-TLR stimulation conditions, but again observed no substantial overlap.

**Figure 3.**
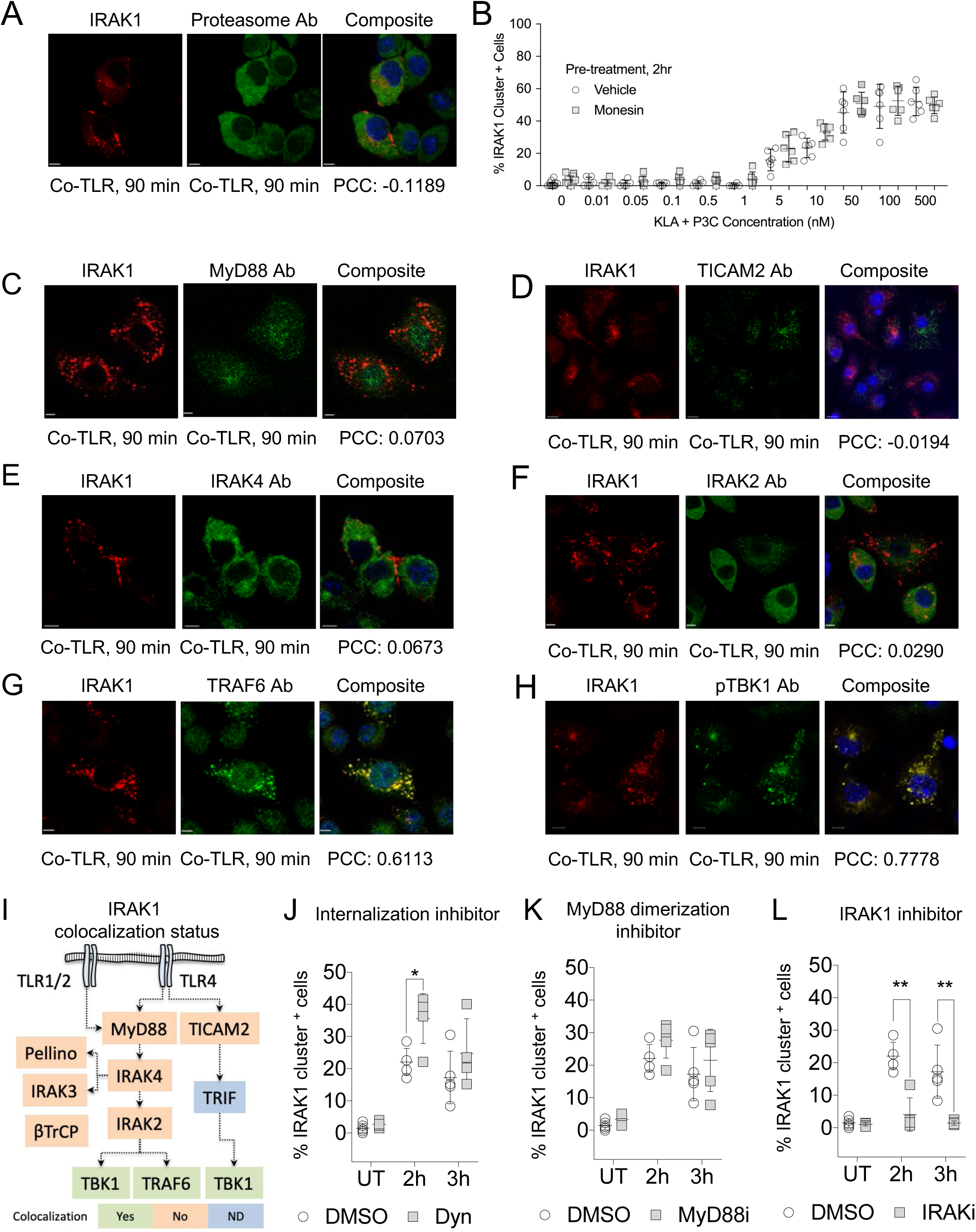
Properties of IRAK1-containing clusters. (A) Co-staining of IRAK1 clusters and proteasome in iBMDMs. Kdo-2 Lipid A and P3C co-treatment (50 nM). Scale Bar: 10*μ*. (B) Dose response of IRAK1 clustering in iBMDMs with Kdo-2 Lipid A and P3C ± 2h pre-treatment with 4uL of monensin, BD GolgiStop™ for every 3 mL of cell culture. (C-K) Kdo-2 Lipid A and P3C co-treatment (50 nM) in iBMDMs. Co-staining of IRAK1 clusters and (C) MyD88, (D) TICAM2, (E) IRAK4, (F) IRAK2, (G) TRAF6 and (H) pTBK1. Scale Bar: 10*μ*. (I) Summary of IRAK1 clustering with TLR signaling components. ND: Not Determined. (J-L) IRAK1 clustering in iBMDMs 0, 2 and 3h post-treatment with 50 nM Kdo-2 Lipid A and P3C ± 30 min pre-treatment with (J) dynasore, an internalization inhibitor (20*μ*M); (K) ST2825, a MyD88 dimerization inhibitor (20*μ*M); and (L) thymoquinone, an IRAK1 kinase inhibitor (25*μ*M). (Panels B, J-L) Data are represented as median ± MAD. (Panels J-L) Unpaired t test with Holm-Šídák’s correction. p = 0.1234 (ns), 0.0332(*); 0.0021 (**); 0.0002 (***); < 0.0001 (****). Data shown are representative of at least two independent experiments. See also supporting Fig. S2.

To determine if IRAK1 clustering was dependent on autocrine, paracrine, or intracrine responses following TLR activation, we pre-treated macrophages with monensin which, through interaction with the Golgi transmembrane Na^+^/H^+^ antiporter, prevents protein secretion from the medial to trans cisternae of the Golgi complex (Mollenhauer et al., 1990). This reduces the cell secretions which would promote autocrine and paracrine signaling. Monensin did not alter the IRAK1 clustering dose response to co-TLR stimulation (Fig. 3B), suggesting that clustering was predominantly due to intracrine signaling events.

Since IRAK1-containing clusters may influence TLR signaling flux under co-TLR stimulation conditions, we used confocal microscopy to analyze the clusters for signaling proteins involved in proximal, medial and distal events downstream of TLR activation. We found that neither the proximal adapter proteins MyD88 or TICAM2 (Fig. 3C, D), nor the medial signaling components IRAK4 (Fig. 3E), IRAK3 (Fig. S2C) or IRAK2 (Fig. 3F) colocalized with co-TLR induced IRAK1 clusters. In contrast, the distal TLR signaling components TRAF6 (Fig. 3G, S2D, E) and pTBK1 (Fig. 3H, S2F, G) showed substantial colocalization with IRAK1 after 90 min of co-TLR stimulation. These observations were recapitulated with proximity ligation assays, which demonstrated that IRAK1-TRAF6 and IRAK1-pTBK1 PLA intensity values were higher in IRAK1 cluster-containing cells and were strongly induced by ligand treatment (Fig. S2H, I). We also observed that IRAK2-TRAF6 and IRAK2-pTBK1 PLA intensities were higher in IRAK1 cluster-negative or *Irak1^−/−^* cells (Fig S2J, K), suggesting a degree of competition between IRAK1 clusters and IRAK2 for distal TLR signaling components. The IRAK1 cluster colocalization status with TLR pathway components at 90 min post-TLR4 and TLR1/2 co-stimulation is summarized in Fig.3I.

Considering the critical role for TRAF6 and TBK1 in TLR signal propagation, we sought to address whether signaling to NF-κB and MAPK was altered under the co-TLR stimulation conditions that promoted localization of TRAF6 and TBK1 to the IRAK1 clusters. We measured nuclear translocation of p-p65 and p-ATF2, as readouts for activity of the NF-κB and MAPK pathways respectively. We observed no substantial difference in the NF-κB or MAPK responses to single or co-TLR activation (Fig. S2L, M). This suggests that co-TLR activated IRAK1 clustering and TRAF6/TBK1 recruitment to these clusters may occur independently of, and/or subsequent to, the canonical signaling through NF-κB and MAPK that drives the TLR-induced transcriptional program. This is supported by the IRAK1-independence of the TLR-induced transcriptional response (Fig. 1B).

To test if co-TLR stimulation induced IRAK1 clustering was dependent on TLR internalization, we used Dynasore to prevent endocytosis through inhibition of dynamin activity and lipid raft organization (Preta et al., 2015). At a concentration of Dynasore that effectively perturbed TLR-induced pATF2 translocation (Fig S2N), IRAK1 clustering was not diminished (Fig. 3J). We also inhibited MyD88 dimerization using ST2825, a peptidomimetic compound that mimics the heptapeptide in the BB-loop of the MyD88-TIR domain thereby interfering with the recruitment of IRAK1 and IRAK4 by MyD88 (Loiarro et al., 2007). ST2825 did not have a detectable effect on co-TLR induced IRAK1 clustering (Fig. 3K) but it did reduce TLR pathway mediated p-ATF2 nuclear translocation (Fig. S2N). This suggested that the reduction in MyD88 dimerization that we could achieve with ST2825, though critical for canonical TLR pathway activity, was not required to form the co-TLR induced IRAK1 clusters. We then asked if IRAK1 kinase activity was necessary for IRAK1 clustering under co-TLR stimulation. We used thymoquinone which is known to inhibit IRAK1 and not IRAK4 kinase activity (Hossen et al., 2017). On pre-treatment with the IRAK1 kinase inhibitor, we observed a marked reduction in co-TLR induced IRAK1 clustering (Fig. 3L). IRAK1 inhibition did not have any effect on TLR activated pATF2 (Fig. S2N), which was consistent with the intact transcriptional response in *Irak1^−/−^* BMDMs (Fig. 1B, (Sun et al., 2016)). This suggested that IRAK1 kinase activity was critical either for the initiation or maintenance of the co-TLR induced IRAK1 clusters but not for the canonical TLR pathway response.

### IRAK1 clusters regulate ASC-containing inflammasomes

Since the peak formation of IRAK1 clusters occurred later than typical TLR-induced NF-κB and MAPK activation (Fig. S1A, D), and we had found the clusters colocalized with the distal TLR signaling components TRAF6 (Fig. 3G), and TBK1 (Fig. 3H), we speculated that the IRAK1 clustering could influence later non-canonical TLR signaling events. Previous studies have demonstrated that under acute inflammasome activation conditions, IRAK1 plays an important role in linking TLR stimulation with rapid NLRP3 activation that does not depend on transcriptional priming of inflammasome components (Fernandes-Alnemri et al., 2013; Lin et al., 2014). It has also been shown that TRAF6 is necessary for transcription independent priming of inflammasomes under single TLR or IL-1R stimulation (Xing et al., 2017). Given these known roles for IRAK1 and TRAF6 in single TLR-induced transcription-independent inflammasome priming, we considered the possibility that the IRAK1 containing clusters observed under co-TLR stimulation may have a role in facilitating inflammasome activation upon multi-PRR stimulation.

We stained for ASC, a key adaptor in nucleation of several classes of inflammasome (Eitel et al., 2010; Franchi et al., 2009; Man and Kanneganti, 2015; Martinon et al., 2002) and observed substantial colocalization with IRAK1 clusters in co-TLR activated macrophages (Fig. 4A, S3A-D). Notably, the discrete ASC clustering pattern induced by co-TLR stimulation was absent in *Irak1^−/−^* macrophages (Fig. 4B). Since co-TLR stimulated iBMDMs gave both IRAK1 and ASC clustering we looked at the correlation between the IRAK1 and ASC clusters across three time points and three pairwise doses of KLA and P3C and saw that the occurrence of IRAK1 and ASC clusters correlated with a correlation coefficient of 0.8302 and p <0.0001 (Fig. 4C).

**Figure 4.**
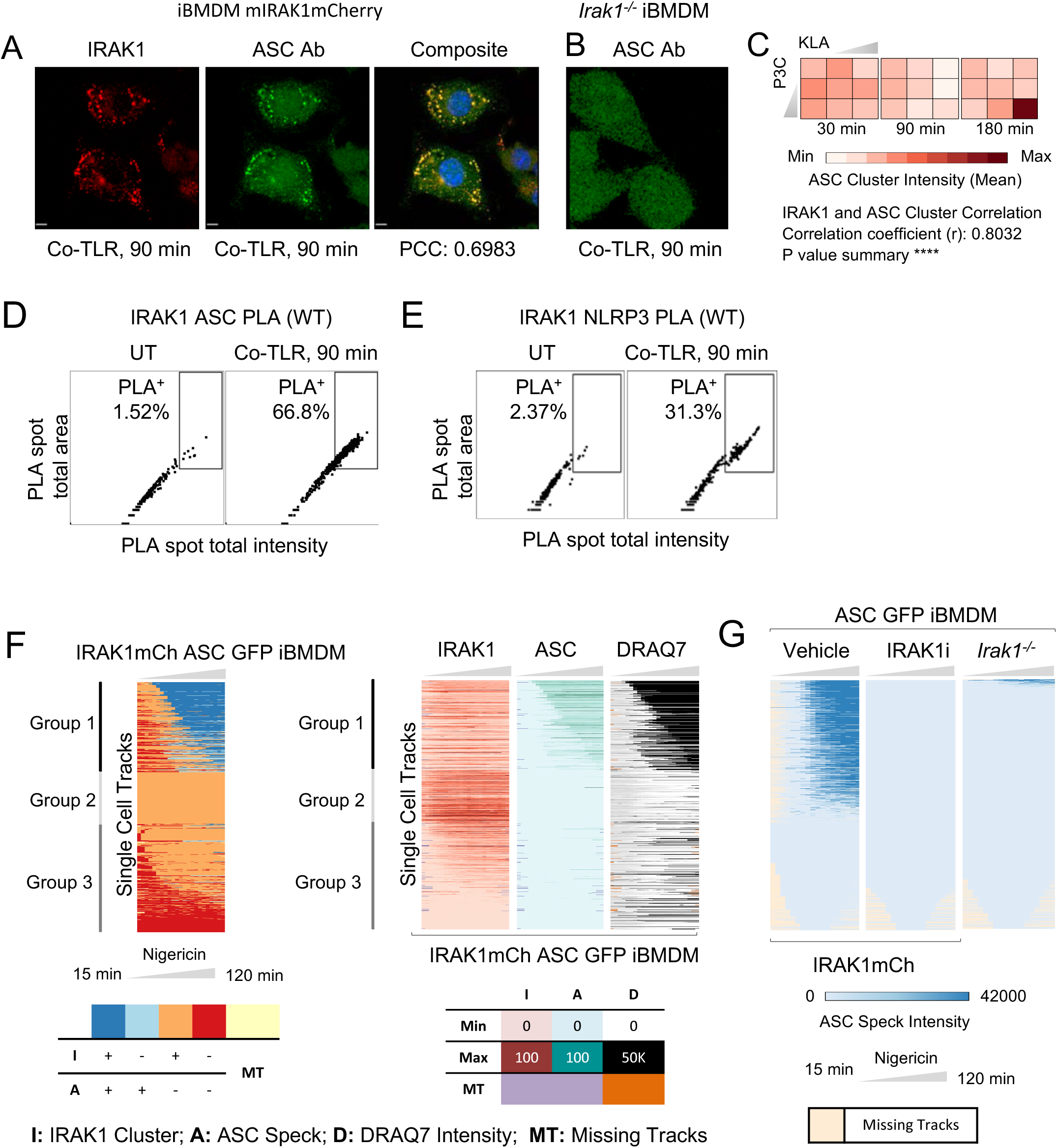
IRAK1 clusters recruit inflammasome components. Kdo-2 Lipid A and P3C co-treatment (50 nM) in iBMDMs. (A) Co-staining of IRAK1 clusters and ASC (SCBT sc-22514-R). Scale Bar: 10*μ*. (B) ASC (SCBT sc-22514-R) staining in *Irak1*^−/−^ iBMDMs. Scale Bar: 10*μ*. (C) Quantification of ASC (SCBT sc-22514-R) clustering in iBMDMs after pairwise stimulation with TLR ligands – Kdo-2 Lipid A (0.5, 50 nM) and P3C (0.5, 50 nM). Data are represented as mean (n = 3). IRAK1 and ASC cluster correlation calculated using IRAK1 clustering data from Fig. 2D. (D, E) Proximity ligation assays in iBMDMs. Data are represented as Median ± SD. (D) IRAK1-ASC PLA, (E) IRAK1-NLRP3 PLA. (F) Single cell live imaging of IRAK1 clustering and ASC clustering in IRAK1mCh ASC GFP iBMDMs on 2h priming with Kdo-2 Lipid A and P3C (50 nM) followed by Nigericin (10 *μ*M) trigger for 2h. The cell tracks could be broadly divided into three groups. Group 1 cells showing ASC specks exhibited IRAK1 clusters for a comparable time period prior to ASC speck formation. The non-ASC specking cells could be divided into two groups - group 2 cells had the largest IRAK1 clusters throughout the time course, while group 3 cells had either very weak or undetectable IRAK1 clusters. (G) ASC specking in IRAK1-mCh ASC-GFP +/− thymoquinone, an IRAK1 kinase inhibitor (25*μ*M) or *Irak1^−/−^* ASC-GFP iBMDMs on 2h priming with Kdo-2 Lipid A and P3C (50 nM) followed by Nigericin (10 *μ*M) trigger for 2h. (Panel C) Paired t test. p = 0.1234 (ns), 0.0332(*); 0.0021 (**); 0.0002 (***); < 0.0001 (****). Data shown are representative of at least two independent experiments. See also supporting Fig. S3, Movies 6-8.

We measured the interaction using PLA and found that IRAK1 and ASC show substantially increased proximity in KLA and P3C co-stimulated macrophages (Fig. 4D). ASC phosphorylation is required for the formation of the speck-like ASC aggregates that form after triggering of an inflammasome response (Hara et al., 2013). Since we showed above that ASC colocalized with IRAK1 clusters, we asked if the IRAK1 cluster-associated ASC is phosphorylated. pASC shows an aggregate pattern in WT iBMDMs (Fig. S3E) which is absent in *Irak1^−/−^* iBMDMs (Fig. S3F) and the pASC aggregates colocalize well with IRAK1 clusters (Fig. S3E, G, H). Additional inflammasome components like NLRP3 were also found in association with IRAK1 clusters upon co-TLR stimulation, although to a slightly lesser degree than ASC (Fig. 4E, S3I, K, L), and some degree of NLRP3 clustering was still observed in *Irak1^−/−^* iBMDMs (Fig. S3J). It has been shown that *Salmonella Typhimurium* induced inflammasomes recruit NLRP3 and NLRC4 to the same macromolecular complex (Man et al., 2014) and that caspase-1 activation by *Pseudomonas aeruginosa* depends on NLRC4 inflammasomes (Franchi et al., 2007; Sutterwala et al., 2007). Since IRAK1 clusters were induced by both of these bacteria (Fig. 2A, B, Movies 1-2), we co-stained for NLRC4 with IRAK1 clusters and observed substantial co-localization (Fig. S3M, O, P). Similar to the observation for NLRP3, some NLRC4 clustering was still evident in *Irak1^−/−^* iBMDMs (Fig. S3N).

Since co-TLR stimulation led to ASC clustering, and ASC speck formation indicates assembly of the inflammasome (Guo et al., 2015; Masumoto et al., 2001; Richards et al., 2001; Stutz et al., 2013), we tested if the co-TLR stimulation induced IRAK1 clustering status of a macrophage affected its propensity to form ASC specks on inflammasome triggering with nigericin. To accomplish this, we generated clones which stably express ASC-GFP at low levels in the IRAK1-mCherry iBMDM cells and also in *Irak1^−/−^* iBMDMs. At these low levels of ASC, there was negligible spontaneous ASC specking in either cell line. We used live cell imaging to track ASC and IRAK1 and observed robust ASC speck formation in a subset of cells in response to co-TLR priming and nigericin triggering (Fig. 4F). In contrast, we detected no ASC specks in cells pre-treated with the IRAK1 inhibitor thymoquinone, and very few ASC specks in the *Irak1^−/−^* iBMDM (Fig. 4G). Taking advantage of the ability to track ASC and IRAK1 in the same single cells, we also noted that the majority of cells showing ASC specks exhibited IRAK1 clusters for a comparable time period prior to ASC speck formation (Fig. 4F, group 1 cell tracks). We also observed that these group 1 cells, exhibiting both ASC specks and IRAK1 clusters, showed the highest DRAQ7 uptake signal (Fig. 4F), an indication of plasma membrane compromise in these cells. Notably, the non-ASC specking cells could be divided into two further groups. Group 2 cells had the largest IRAK1 clusters throughout the time course and the lowest DRAQ7 signal, suggesting that IRAK1 clustering by itself did not diminish cell viability, while group 3 cells had either very weak or undetectable IRAK1 clusters (Fig. 4F). These data suggest that there may be an optimal intermediate magnitude for the IRAK1 clusters to support ASC activation.

### IRAK1 clusters facilitate licensing of inflammasomes in multi-PRR primed macrophages

Since ASC and other inflammasome components colocalized with co-TLR induced IRAK1 clusters in iBMDMs, we asked if a similar pattern was evident in primary macrophages. PLA assays confirmed proximity of ASC (Fig. 5A, B), pASC (Fig. S4A, B) and NLRP3 (Fig. 5C, D) with IRAK1 in BMDM challenged with either co-TLR ligands or heat killed bacteria. We further tested if this persisted following an inflammasome trigger and observed proximity of IRAK1 with ASC, pASC and NLRPs in response to nigericin treatment under multiple priming conditions (Fig. S4C-E).

**Figure 5.**
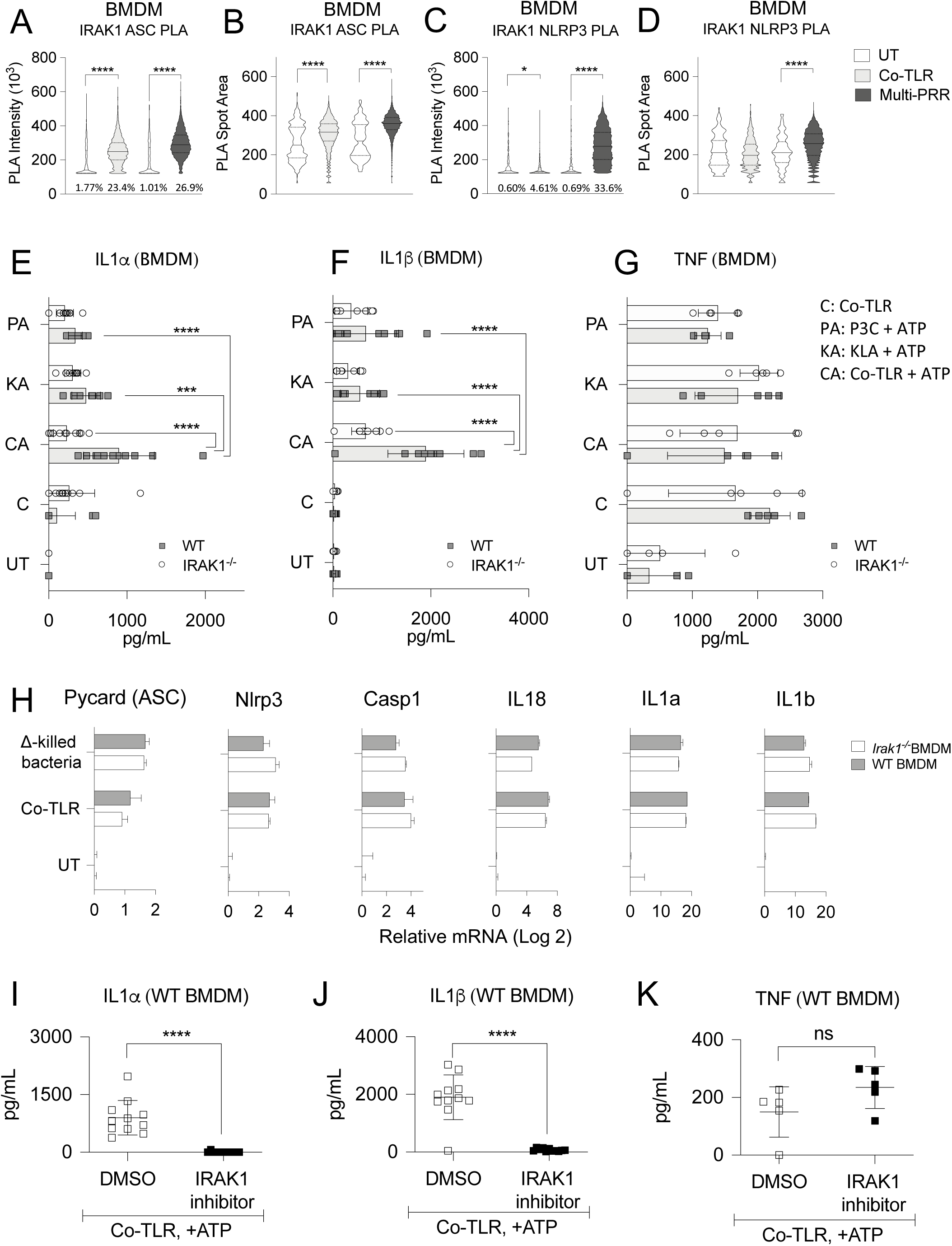
IRAK1 clusters are required for licensing of the inflammasome in BMDMs. (A-D) Proximity ligation assay in BMDMs. At least 20,000 cells imaged for all graphs. Percentage of cells showing PLA spots stated at the base of the graphs. Co-TLR stimulation with Kdo-2 Lipid A and P3C co-treatment (500 nM each). Multi-PRR stimulation using heat-killed *Yersinia pseudotuberculosis* (MOI 100). (A) IRAK1-ASC PLA intensity, (B) IRAK1-ASC PLA spot area, (C) IRAK1-NLRP3 PLA intensity, (D) IRAK1-NLRP3 PLA spot area. (E-G) Cytokine secretion measured by ELISA in BMDMs primed with single TLR ligands: Kdo-2 Lipid A or P3C (1 uM each) or with co-TLR stimulation using 500 nM of both ligands, followed by ATP (5 mM) trigger for 2h. Mean ± SD from at least 4 wells. (E) IL1*α*, (F) IL1*β* and (G) TNF. (H) qPCR quantification of *Pycard (ASC)*, *Nlrp3*, *Casp1*, *IL18*, *IL1a* and *IL1b* in BMDMs primed with heat-killed *Yersinia pseudotuberculosis* (MOI 100) or with co-TLR stimulation using 500 nM of both Kdo-2 Lipid A and P3C. Mean ± SD from at least 4 wells. (I-K) Cytokine secretion measured as detailed in E-G in BMDMs pretreated with thymoquinone, an IRAK1 kinase inhibitor (25*μ*M). (I) IL1*α*, (J) IL1*β* and (K) TNF. (Panels A-D) Kolmogorov-Smirnov test. (Panels E-G, K-M) Tukey’s multiple comparisons test. (Panels H-J) Two-way ANOVA with Geisser-Greenhouse correction and Šídák’s multiple comparison test. p = 0.1234 (ns), 0.0332(*); 0.0021 (**); 0.0002 (***); < 0.0001 (****). See also supporting Fig. S4.

Given the colocalization of IRAK1 clusters with inflammasome components in co-PRR or co-TLR primed cells prior to triggering (Fig. 4A,C-F, S3, 5A-D, S4A, B), we tested if co-TLR priming leads to a more robust inflammasome response compared to single TLR ligand priming. To account for the higher ligand dose in the co-TLR primed condition, the single ligand treatments used twice the concentration employed in co-stimulation conditions. In co-TLR primed cells, we observed substantially higher IL-1α (Fig. 5E) and IL-1β (Fig. 5F) release compared to cell primed with single TLR ligands. Notably, we found that all measured IL-1 outputs were significantly attenuated in *Irak1^−/−^* BMDM only on co-TLR priming but not with single-TLR priming (Fig. 5E, F). TNF levels, which represent a surrogate for strength of priming, were not substantially different between WT and *Irak1^−/−^* BMDMs nor between single or co-TLR activated cells (Fig. 5G).

To determine whether the effect of IRAK1 on the inflammasome response was related to the efficacy of TLR-induced transcriptional priming of critical inflammasome components, we measured induction of mRNA for *Pycard (ASC), Nlrp3, Casp1,Il18, IL1a and Il1b* in response to either co-TLR or heat-killed bacteria stimulation in WT and *Irak1^−/−^* BMDM (Fig. 5H). We observed no significant difference in transcriptional priming of these critical inflammasome genes, suggesting that IRAK1 does not play a major role in TLR-induced transcriptional priming and also consistent with our previously observed signaling and transcription responses in *Irak1*^−/−^ cells (Fig. 1B; (Sun et al., 2016)). Moreover, the enhanced IL-1α and IL-1β release in co-TLR treated cells (Fig. 5E, F) is not simply a reflection of the magnitude of Il1a and Il1b mRNA induction compared to single TLR activated cells (Fig. S4F, G), and the IL-1 gene mRNA levels are similarly unaffected by IRAK1 deficiency. This again distinguishes the role of IRAK1 in inflammasome activation from the transcriptional priming function that might be expected of a TLR pathway component and suggests a more direct role for the IRAK1 protein in facilitating inflammasome function.

IRAK1 kinase inhibition with thymoquinone in WT BMDMs phenocopied *Irak1*^−/−^ BMDMs in reducing trigger induced IL-1α and IL-1β release, while not affecting the canonical TLR pathway induction of TNF (Fig. 5I-K). This suggested that either IRAK1 auto-phosphorylation or IRAK1 phosphorylation of cluster components could be involved in either initiating or stabilizing IRAK1 clusters. Post translational modifications (PTMs) of inflammasome components are known to be involved in licensing of inflammasomes (Hoss et al., 2017; Yang et al., 2017). We have presented multiple lines of evidence (Fig. 1B, E, 4G, 5E-K, S4F,G) suggesting that IRAK1 is not required for TLR-activated transcriptional priming of inflammasome components but is critical for inflammasome activation in multi-PRR primed cells. We therefore investigated whether IRAK1, and/or components of the IRAK1 cluster complex, could be involved in regulating PTMs required for inflammasome licensing.

### IRAK1 clusters are dependent on a JNK MAPK cascade

MAPKs have been proposed to regulate inflammasome activity and it has been shown that JNK mediated phosphorylation of ASC can act as a molecular switch, controlling its ability to form specks and support inflammasome assembly (Hara et al., 2013). To establish if co-TLR induced IRAK1 clusters were regulated by JNK, we measured IRAK1 clustering in the presence of a JNK inhibitor and observed a strong reduction in cluster frequency (Fig. 6A). pJNK1/2 also formed aggregates and colocalized with co-TLR induced IRAK1 clusters in iBMDMs (Fig. 6B, S5A, B), while these pJNK1/2 aggregates were absent in *Irak1^−/−^* iBMDMs (Fig.S5C). PLA assays demonstrated increased IRAK1 and pJNK proximity in WT BMDMs that were primed with either multi-PRR or co-TLR stimulation (Fig. 6C, D), and this persisted after nigericin triggering (Fig. S5D). JNK inhibition in WT BMDMs phenocopied *Irak1^−/−^* BMDMs in reducing inflammasome-mediated IL-1α and IL-1β release (Fig. 6E, F) with no substantial effect on TLR-induced TNF secretion (Fig. 6G).

Since the MAPK kinase MKK7 has been implicated in regulation of inflammation-mediated JNK activation (Tournier et al., 2001), we imaged pMKK7 under IRAK1 clustering conditions and observed strong pMKK7 colocalization with IRAK1 clusters (Fig. 6H, S5E, F), and a lack of MKK7 clustering in *Irak1^−/−^* iBMDMs (Fig. S5G). Similar to that observed for pJNK, the pMKK7-IRAK1 proximity assessed by PLA was sustained after nigericin treatment (Fig. S5H). In contrast to pMKK7, pMKK4, a MAPK kinase that activates both p38 MAPK and JNK (Haeusgen et al., 2011) does not colocalize with the IRAK1 clusters as well as pMKK7 (Fig. S5I).

**Figure 6.**
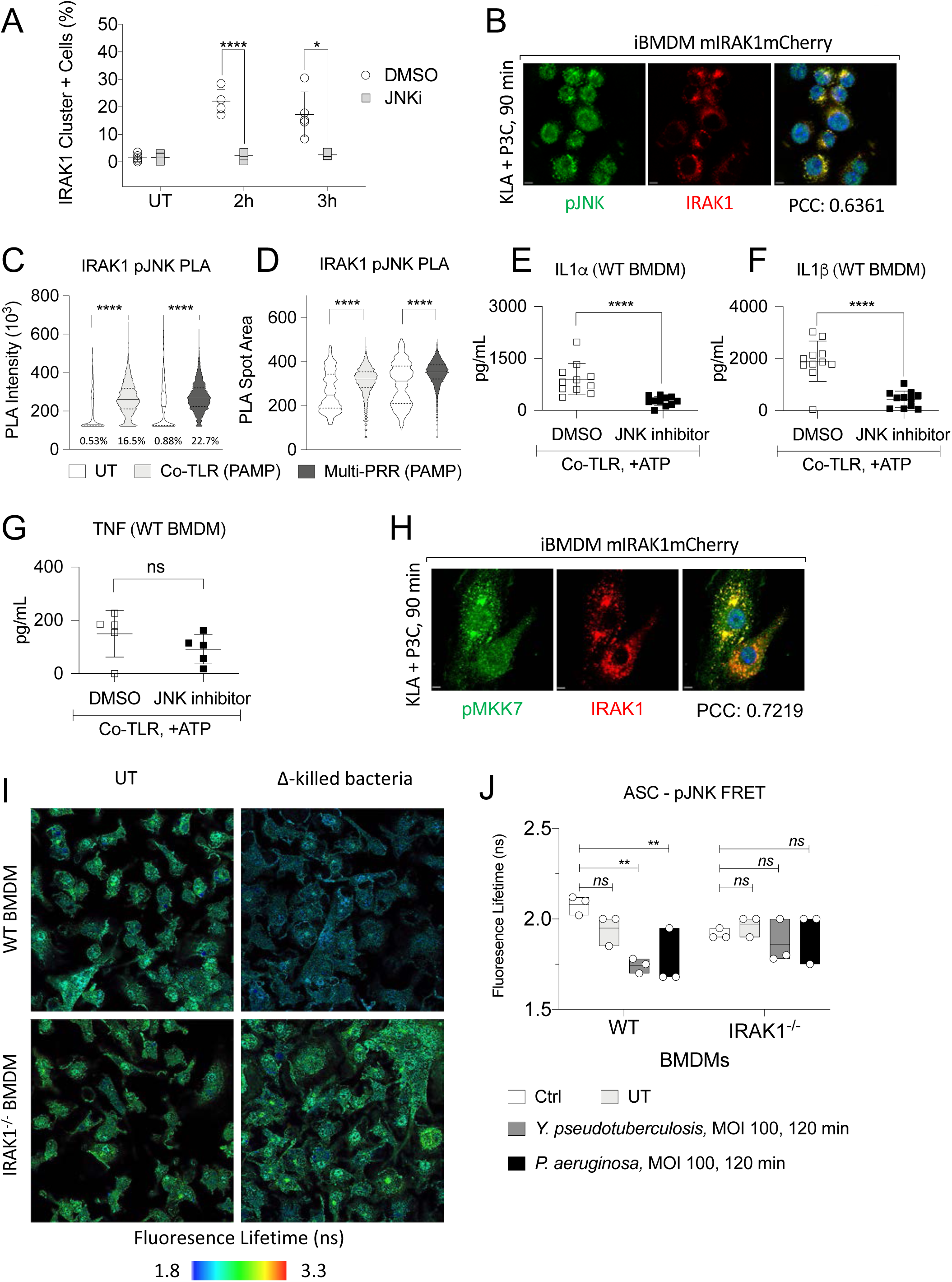
IRAK1 clusters are dependent on a recruited JNK MAPK cascade. (A) IRAK1 clustering in iBMDMs 0, 2 and 3h post-treatment with 50 nM Kdo-2Lipid A and P3C ± 30 min pre-treatment with JNK inhibitor (10 *μ*M). (B) Co-staining of IRAK1 clusters and pJNK in iBMDMs. Kdo-2 Lipid A and P3C co-treatment (50 nM). Scale Bar: 10*μ*. (C, D) Proximity ligation assay in BMDMs. At least 20,000 cells imaged for all graphs. Co-TLR stimulation with Kdo-2 Lipid A and P3C co-treatment (500 nM). (C) IRAK1-pJNK PLA intensity, (D) IRAK1-pJNK PLA spot area. (E-G) Cytokine secretion measured by ELISA in BMDMs pretreated with JNK inhibitor, 10*μ*M, primed with single TLR ligands: Kdo-2 Lipid A or P3C (1 uM each) or with co-TLR stimulation using 500 nM of both ligands, followed by ATP (5 mM) trigger for 2h. (E) IL1*α*, (F) IL1*β* and (G) TNF. (H) Co-staining of IRAK1 clusters and pMKK7 in iBMDMs. Kdo-2 Lipid A and P3C co-treatment (50 nM). Scale Bar: 10*μ*. (I,J) Fluorescence lifetime measurement of ASC where the acceptor/donor pair was pJNK/ASC primed with multi-PRR stimulation by heat-killed *Yersinia pseudotuberculosis* or *Pseudomonas aeruginosa* (MOI 100) in WT and *Irak1*^−/−^ BMDMs. (Panel A) Unpaired t test with Holm-Šídák’s correction. p = 0.1234 (ns), 0.0332(*); 0.0021 (**); 0.0002 (***); < 0.0001 (****). (Panels C,D) Kolmogorov-Smirnov test. (Panels E-G) Tukey’s multiple comparisons test. (Panel J) Two-way ANOVA and Šídák’s multiple comparison test. p = 0.1234 (ns), 0.0332(*); 0.0021 (**); 0.0002 (***); < 0.0001 (****). Data shown are representative of at least two independent experiments. See also supporting Fig. S5.

Given the IL-1 response defects and increased mortality in *Irak1^−/−^* mice infected with *Pseudomonas aeruginosa* or *Yersinia pseudotuberculosis* respectively, we tested whether IRAK1 influenced ASC and JNK interaction after bacterial challenge. WT and *Irak1^−/−^* BMDMs were primed with heat-killed *Yersinia pseudotuberculosis* or *Pseudomonas aeruginosa* for 2 hours and the cells were fixed. ASC-pJNK interaction was measured by FRET/FLIM imaging where increased ASC-pJNK interaction would lead to a reduction in the fluorescence lifetime (FLIM) (Fig. 6I, J). Significant FLIM decreases were observed in WT BMDM challenged with either bacteria while no significant decrease was evident in *Irak1^−/−^* BMDMs (Fig. 6I, J). Taken together, these observations implicate IRAK1 in licensing the ASC inflammasome through a JNK-dependent process when macrophages are faced with a complex priming stimulus.

Immune signaling is known to proceed through cooperative assembly of supra molecular organizing complexes (SMOCs) (Algeciras-Schimnich et al., 2002; Bryant et al., 2015; Hou et al., 2011; Kagan et al., 2014; Lin et al., 2010; Lu et al., 2014; Motshwene et al., 2009; Park et al., 2007; Qiao and Wu, 2015; Vajjhala et al., 2017). Minimal SMOC properties are that they are 1) multimeric clusters of proteins; 2) known to regulate and organize signaling; 3) inducible upon specific stimulation; and 4) have a distinct cellular location. We have provided evidence that the clustering of IRAK1 under conditions of co-TLR or multi-PRR stimulation meets all of these criteria, suggesting that these IRAK1 clusters may be a previously unrecognized innate immune signaling SMOC. They are also distinct from the known SMOCs in that these IRAK1 coincidence detection clusters include adaptors from multiple signaling pathways, namely; TLR, MAPK and inflammasome pathways.

## Discussion

Our knowledge of innate immune responses to multi-PRR stimulation and regulation of PRR pathway crosstalk is limited, but this understanding is critical since outside of laboratory settings, macrophages rarely encounter single PAMP or DAMP stimuli. Collaboration of multiple PRRs and their adaptors is required for an optimal host response to many pathogens (Delaloye et al., 2009; Ferwerda et al., 2007; Slater et al., 2010). In the context of vaccine-adjuvant development, it has also been shown that co-delivery of TLR agonists as adjuvants elicits an improved immune memory response (Ebrahimian et al., 2017; Pulendran and Ahmed, 2011). Thus, a cogent and effective immune response must involve concomitant engagement of multiple PRR pathways. Few prior studies have identified specific PRR signaling events that are selectively induced by multi-ligand treatment. We demonstrate a discrete IRAK1 clustering response that occurs in macrophages challenged with either combined TLR ligands or intact bacteria.

While there is emerging evidence that the inflammasome priming signals from TLR activation of macrophages go beyond mRNA elevation of inflammasome components, this is difficult to prove genetically as knock-out of the TLR components that are critical for the priming step will prevent the required mRNA induction. However, the redundant functions of IRAKs in mouse cells (Kawagoe et al., 2008; Sun et al., 2016; Swantek et al., 2000) have allowed us to uncover a previously unappreciated aspect of TLR/inflammasome crosstalk, as *Irak1^−/−^* has no effect on mRNA priming of inflammasome genes. Notably, *Irak1^−/−^* also has no effect on the overall process of inflammasome activation when cells are primed with single TLR ligands, which may explain why the function of IRAK1 that we describe here has not been identified in prior studies. While the transcriptional response of *Irak1^−/−^* BMDMs was similar to WT BMDMs, the ability of ASC, NLRP3, JNK, pMKK7, TRAF6, and pTBK1 to form clusters in response to multi-PRR stimulation was contingent on the presence of IRAK1. This allowed us to demarcate that the formation of an IRAK1 SMOC that links the valency of TLR signaling with inflammasome licensing can occur independently of the canonical TLR signaling and transcription response.

Since the initial inflammasome priming stimulus in vivo is more likely to involve multiple PAMPs or DAMPs, we believe that the IRAK1 SMOC-dependent licensing would occur in most physiological scenarios that lead to inflammasome activation, and thus explain the susceptibility of *Irak1^−/−^* mice to infections where the IL-1 family cytokines have a critical host protective role. It is also possible that the IRAK1-dependent licensing function contributes to innate immune ‘threat assessment’ as recently described (Evavold and Kagan, 2019), as IRAK1-SMOCs are more strongly induced by bacteria than purified ligands.

IRAK1 has been shown to propagate early signaling after TLR activation, while IRAK2 is important to sustain the response (Kawagoe et al., 2008). This model fits with our observation that IRAK1 clusters form in the time period subsequent to initial TLR signaling events. While proximal TLR signaling components such as TRAF6 and TBK1 are recruited to the IRAK1 SMOC, we would propose that their function in propagating the canonical TLR signal is already fulfilled at this point. Local concentrations of IRAK1 are bound to increase due to IRAK1 clustering and consequently, components that interact with the IRAK1 cluster could signal more efficiently. This process is known as signaling flux redistribution and has been proposed as a way TLR signaling could be regulated (Selvarajoo et al., 2008). IRAK1 has three TRAF6 binding sites while IRAK2 has two sites (Gosu et al., 2012). This difference in TRAF6 binding capacity would be amplified by IRAK1 clustering and could be a means by which IRAK1 competes with IRAK2 for distal TLR signaling components.

The recruitment of both TRAF6 and TBK1 to the IRAK1 SMOC is also noteworthy from a metabolic perspective, as it has been recently reported that these TLR pathway components are activated downstream of the myddosome to induce the glycolytic shift observed in inflammatory macrophages (Tan and Kagan, 2019). It is possible therefore that IRAK1 SMOCs participate in the induction of metabolic changes required to support the energetic requirements of inflammasome activation. This may also contribute towards additional key mitochondrial events that have been implicated in inflammasome activation, such as localization of inflammasome components to the mitochondrial membrane and release of mitochondrial stress signals as inflammasome triggers.

Myddosomes are a principal class of SMOCs in TLR signaling that are known to contain IRAKs. However, the multi-PRR-induced IRAK1 SMOC does not include components of the myddosome and its formation seems to be insensitive to the levels of MyD88 dimerization. In zebrafish, IRAK1 has been shown to regulate a PIDDosome SMOC that drives tumor resistance to radiation therapy independent of MyD88, thereby linking the DNA damage response pathway with TLR signaling (Liu et al., 2019). In fibroblasts, IRAK1 forms clusters in response to high doses of IL-1β, acting as a dose-sensing negative feedback node responsible for limiting signal flow in strongly activated cells (DeFelice et al., 2019). Thus, there are reported examples of IRAK1 acting as a signal regulating hub that can integrate different signaling pathways and sense the magnitude of input stimuli.

Preliminary testing of other TLR pairs showed IRAK1 SMOCs could be formed with other TLR pairs but not as efficiently as with TLR4 and TLR2 co-stimulation. These SMOCs could also be formed albeit to a smaller size when stimulated with complex DAMPs like biglycan that are known to engage TLR4 and TLR2. Further investigations are required to determine if IRAK1 SMOCs formed with different TLR pairs also engage MAPK and inflammasome pathways, or if different ligand combinations push IRAK1 SMOCs to engage alternate effectors.

LPS is a potent PAMP, and modifications of the LPS structure are prevalent in bacteria. It remains to be seen if the different variants of LPS induce IRAK1 SMOC formation on co-stimulation with TLR2 or with other TLRs. Additionally, we do not know how co-stimulation of TLRs leads to assembly of an IRAK1 SMOC. Since the IRAK1 SMOC formation begins to occur within 30 min of ligand activation, it is less likely that cytosolic PRRs and ligand-induced transcriptional events are involved in its initiation. Instead, IRAK1 SMOC formation may require distinct post translational modifications in IRAK1 or other effectors preferentially induced by TLR ligand pairs.

The priming and triggering steps of inflammasome activation are well established, however it is likely that post-translation modifications (PTMs) of inflammasome proteins play an important role in licensing this critical inflammatory process. The lack of a specific identified ligand for NLRP3 might suggest that disparate cellular stresses can induce signaling events that coordinate the PTMs of inflammasome components required to trigger a response. Myddosomes and inflammasomes are separate SMOCs induced during priming and triggering respectively, but there is no prior evidence that components typically associated with one or the other become associated during the licensing phase. We propose that IRAK1 recruits a MKK7/JNK complex to facilitate licensing of key inflammasome components, with ASC being the most likely substrate based on previous work (Hara et al., 2013). We observe a dependence on both IRAK1 and JNK kinase activity for the inflammasome response in co-TLR or multi-PRR primed macrophages, and propose a model whereby JNK kinase activity is required to sustain IRAK1 clusters and IRAK1 kinase activity supports JNK-ASC proximity and ASC speck formation. Live cell imaging of IRAK1 clusters and nigericin-triggered ASC specks also suggests a consistent time duration between onset of IRAK1 clustering and ASC speck formation, and that an intermediate intensity of IRAK1 clusters correlates directly with ASC speck-positive cells. This implies that there may be a quantitatively optimal window for the upstream protein modification events that license inflammasome formation, and that in a given population of cells, only a subset will meet this requirement. It remains to be seen whether this represents some sort of checkpoint for cellular fitness to mount an inflammatory response.

Our data suggest that the role of TLR activation in inflammasome activation goes beyond the transcriptional priming of inflammasome mRNAs, and that proteins considered central to TLR pathway activation play additional roles in facilitating the inflammasome response when innate immune cells are faced with complex microbial and danger signals. Elucidating how these pathway crosstalk events are integrated and regulated in both healthy and disease states will be an important focus of future research.

## Acknowledgements

This work was supported by the Intramural Research Program of NIAID, NIH. *Irak1*^−/−^ mice were provided by Dr.Chadrashekhar Pasare, Cincinnati Children’s Hospital Medical Center. GFP expressing *Salmonella Typhimurium* were provided by Dr.Edward A. Miao, University of North Carolina at Chapel Hill School of Medicine. We acknowledge Dr. Andrea J. Radtke, Lymphocyte Biology Section, Laboratory of Immune System Biology, NIAID for initial training with FlowJo^®^ based analysis of immunofluorescence data. We thank Drs. Sesha Tekur and Nicholas Radio from Thermo Fisher Scientific for assistance with the initial optimization of the spot detection algorithm on the CellInsight NXT imager.

## Author Contributions

Conceptualization, S.J.V., O.E., C.J.B., and I.D.C.F.; Methodology, S.J.V., M.S., O.E., J.S., C.J.B., M.G.D., J.L., N.B., R.A.G., K-S.O., G.,P., S.G., R.R.V. and I.D.C.F.; Software, S.J.V., R.J.C., S.G. and G.P.; Validation, S.J.V., M.S., O.E., C.J.B. and J.L. Formal Analysis, S.J.V., G.P., and I.D.C.F.; Investigation, S.J.V., M.S., O.E., J.S., C.J.B., M.G.D., J.L., N.B., R.A.G., K-S.O., S.G. and G.,P; Resources, G.P., D.D.N., E.L., Y.B., R.R.V. and I.D.C.F.; Data Curation, S.J.V., O.E., J.S., C.J.B., M.G.D., N.B., S.G., G.P., and I.D.C.F.; Writing – Original Draft, S.J.V. and I.D.C.F.; Writing – Review & Editing, S.J.V., M.S., O.E., R.J.C., J.S., C.J.B., M.G.D., N.B., G.P., D.D.N., R.R.V. and I.D.C.F.; Visualization, S.J.V., O.E., C.J.B., G.P., S.G. and I.D.C.F., Supervision, M.S., S.G., G.P., Y.B., R.R.V., and I.D.C.F.; Project Administration, S.J.V. and I.D.C.F.; Funding Acquisition, I.D.C.F.

## Methods

### Generation of iBMDM cell lines

#### mIRAK1mCherry Irak1^−/−^ iBMDMs

The pR-IRAK1-mCherry retroviral plasmid, expressing murine IRAK1 with a C-terminal fused mCherry, was generated by amplifying IRAK1 from an expression vector with the specific primers: forward 5’-TTTGGATCCATGGCCGGGGGGCCGG-3’; and reverse 5’-AAACTCGAGGCTCTGGAATTCATCACTTTCTTCAGGTC-3’. The resultant PCR product was digested with BamHI and XhoI and cloned in-frame into the corresponding sites of a pR-mCherry retroviral vector. Irak1^−/−^ immortalized-bone marrow derived-macrophages (iBMDMs) expressing murine IRAK1-mCherry were generated by retroviral transduction with pR-IRAK1-mCherry using a previously described protocol (Cardona Gloria et al., 2018). FACS sorting for mCherry positive cells was subsequently performed to enrich for IRAK1-mCherry expression.

#### ASC-GFP expressing iBMDMs

Lentiviral plasmid pLEX-MCS-ASC-GFP (Addgene #73957) and packaging plasmids pCMV-VSV-G (Addgene #8454) and pCMV-delta-R8.2 (Addgene #12263) were transfected into adherent 293T17 cells using the TransIT-Lenti transfection system (Mirus). After 72 hours, supernatant was collected, and virus was concentrated using Lenti-X Concentrator (Takara). Existing mIRAK1mCherry and Irak1^−/−^ iBMDM were then transduced with concentrated lentivirus. After 72 hours of transduction, lentivirus-containing media was removed, and puromycin-containing media was added to select for transduced cells over 10 days. The resulting selected populations were then subjected to limiting dilution to isolate monoclonal iBMDM mIRAK1mCherry Irak1^−/−^ASC-GFP and iBMDM Irak1^−/−^ASC-GFP cell lines. Clones used in this study were selected based on moderate ASC-GFP fluorescent signal and lack of ASC speck formation in the absence of priming and triggering stimuli.

### Cell culture and stimulation with TLR ligands

Immortalized Murine Bone Marrow Derived Macrophages (iBMDMs), derived from wild type and *Irak1^−/−^* mice (Swantek et al., 2000), were maintained in Dulbecco’s modified Eagle medium (Glucose, 4.5 g/L), 10% fetal bovine serum (FBS), 20 mM Hepes, and 2 mM glutamine. Bone Marrow Derived Macrophages (BMDMs) from wild type and *Irak1^−/−^* mice were prepared by differentiation for 6 days in the same culture medium containing macrophage colony-stimulating factor (M-CSF; 60 ng/mL, R&D Systems). LPS was from Alexis Biochemicals (Salmonella minnesota R595 TLR grade, catalog no. ALX-581-008-L002), P3C was from InvivoGen (catalog no. tlrl-pms), R848 was from InvivoGen (catalog no. tlrl-r848), and Kdo2-Lipid A P2C was from Avanti Polar Lipids (catalog no. 699500).

### Mouse infections and LPS challenge

All mice were maintained in specific pathogen-free conditions, and all procedures were approved by the National Institute of Allergy and Infectious Diseases Animal Care and Use Committee (National Institutes of Health, Bethesda, MD).

#### Yersinia pseudotuberculosis infection

For infection, *Y. pseudotuberculosis* (strain 32777) was grown in 2XYT media (Quality Biological) overnight at 25°C with vigorous shaking. WT and *Irak1^−/−^* mice (Swantek et al., 2000) were maintained on a C57BL/6J background and were bred from heterozygous *Irak1−/−* mice to generate WT and KO littermates for infection studies. Mice were fasted for 12 hr prior to infection with 1 × 10^7^ CFU bacteria via oral gavage, and at 3- and 5-days post infection were bled retro-orbitally and serum was collected. Serum cytokine levels were quantified by CBA assay.

#### Pseudomonas aeruginosa infection

WT and *Irak1^−/−^* mice at 8 to 12 weeks old were anaesthetized with isoflurane before being inoculated intra-nasally with 50µL of *Pseudomonas aeruginosa strain PA01* diluted to 1×10^9^ CFU/mL in PBS. Control mice were given 50 µL of PBS. Mice were allowed to wake up and were placed in boxes with food and water for 24 hours. At this time point, mice were euthanized by CO_2_ inhalation. Broncho-alveolar lavage (BAL) was performed by flushing the lungs three times using 1mL of ice-cold PBS + 2mM EDTA. BALs were centrifuged at 500g for 5 min to separate cells and supernatant. Protein levels in supernatants were quantified by CBA assay.

#### Endotoxic shock

WT and *Irak1^−/−^* mice were injected intraperitoneally with 10mg per kg of body weight lipopolysaccharide from *Salmonella enterica* serotype Minnesota (Sigma) dissolved at 1mg/mL in sterile PBS. Mice were weighed twice per day for up to 10 days, after which any surviving mice were euthanized by CO_2_ inhalation. Survival curves were analyzed using the log-rank (Mantel-Cox) test.

### High-content imaging of iBMDMs and BMDMs

Macrophages seeded in 384 well plates plastic-bottom (iBMDMs) (Falcon, 353962) and glass-bottom (day-6 BMDMs) (MatTek, PBK384G-1.5-C) were prepared as explained below. Briefly the iBMDMs and day 6 BMDMs were seeded 24 hours before they were treated with TLR ligands. For imaging ASC specks and ASC PLAs, the iBMDMs and BMDMs were treated with the appropriate TLR ligands followed by a nigericin (10 *μ*M) trigger. Post stimulation, the cells were fixed by incubating with 4% paraformaldehyde for 20 minutes, followed by blocking and permeabilization with PBST-BSA (5% [w/v] bovine serum albumin [BSA] in 0.1% [v/v] Tween 20 in 1× phosphate-buffered saline [PBS]). The fixed cells were treated with the relevant primary antibody at 4°C overnight. Following overnight incubation, the cells were washed three to five times in PBST-BSA. These cells were then incubated for at least 1 h at room temperature with the appropriate secondary antibody. This was followed by PBST-BSA wash (three to five times). Cells were imaged with Cell Insight NXT (ThermoFisher) at 20X, Cell Insight CX7 (ThermoFisher) at 40X, or Opera Phenix High Content Screening System (Perkin Elmer) at 63X. Image analysis on the NXT and CX7 was done using the HCS Studio (ThermoFisher) General Intensity Tools. Image analysis on the Opera images was done using an R analysis pipeline. For spot counting, nuclei were segmented based on the Hoechst 33342 (ThermoFisher, catalog no. R37605) channel, followed by dilating a ring region around the nucleus and detecting the spots in the ring region. Formation of the spots was measured as an increase in the mean integrated spot intensity in the relevant channel.

### Confocal imaging of macrophages

iBMDMs and day-6 BMDMs were prepared as described above. BMDMs for IRAK1 immunofluorescence analysis were permeabilized with cold methanol for 20 minutes followed by rehydration in PBS for 30 min. These cells were imaged on SP8 (Leica) at 63X. The images were analyzed on Imaris 9.2 (Bitplane). Spot counting was done using surface object generation in the relevant channel.

### Live cell imaging of macrophages

mIRAK1mCherry ASC-GFP *Irak1^−/−^* iBMDMs seeded on 96-well plates were imaged at 20X on the CellInsight CX7 with an onstage incubator for ASC speck and IRAK1 cluster counting. The cells were scored for ASC speck and IRAK1 cluster status at each time point and the single live cell tracks were sorted based on a weighting system as follows: ASC speck^+^ IRAK1 cluster^+^ > ASC speck^+^ IRAK1 cluster^−^ > ASC speck^−^ IRAK1 cluster^+^ > ASC speck^−^ IRAK1 cluster^−^. Live cell tracks that lost track of cells for more than 3 time points in the time course were discarded from the analysis. mIRAK1mCherry *Irak1^−/−^* iBMDMs seeded on 8-chamber coverglass slides were infected with GFP expressing *Salmonella Typhimurium* and imaged at 63X on SP8 (Leica). mIRAK1mCherry *Irak1^−/−^* iBMDMs seeded on Bioptechs interchangeable coverglass dish were treated with TLR ligands and imaged at 60X on optical components built around an Olympus IX71 fluorescence. For wide field illumination, the microscope was connected to a metal halide Lambda-XL light source and an excitation filter wheel (Sutter Instruments) fitted with excitation filters.

### Colocalization analysis

We used Pearson’s correlation coefficient (PCC) and Mander’s colocalization coefficient (MCC) as calculated by Imaris 9.2 (Bitplane) software as a statistic for quantifying colocalization. Intensity correlation analysis (ICA) shows that staining intensities of proteins from the same structure vary synchronously, while if they are on different structures the variation will be asynchronous. ICA was run using JACoP for the colocalization images.

### Proximity ligation assay in macrophages

Macrophages seeded in 384 well plates plastic-bottom (iBMDMs) (Falcon, 353962) and glass-bottom (day-6 BMDMs) (MatTek, PBK384G-1.5-C) were prepared as explained above in the methods for high-content imaging of iBMDMs and BMDMs. Post antibody treatment the cells were treated with Duolink PLA reagents according to the manufacturer’s instructions (DUO92002, DUO92004 or DUO92006, DUO82049 and DUO92014; MilliporeSigma). These cells were then imaged on the Cell Insight NXT (ThermoFisher) at 20X, Cell Insight CX7 (ThermoFisher) at 40X and SP8 (Leica) at 100X. PLA spots were selected for analysis only if they were larger and brighter than the following controls: Reaction pairs that lacked one of the primary antibodies or secondary antibodies; Reaction pairs that replaced one of the primary antibodies with the appropriate antibody isotype that matched the missing primary antibody; Reaction pairs that lacked both primary antibodies.

Antibody pairs used for PLA in iBMDMs: IRAK1 Ms (SCBT sc-5288) and TRAF6 Rb (Bioss bs-1184R); IRAK1 Ms (SCBT sc-5288) and pTBK1 Rb (CST 5483S); IRAK1 Ms (SCBT sc-5288) and pTAB2 Rb (CST 8155S); IRAK2 Ms (SCBT sc-515885) and TRAF6 Rb (Bioss bs-1184R); IRAK2 Ms (SCBT sc-515885) and pTBK1 Rb (CST 5483S); IRAK2 Ms (SCBT sc-515885) and pTAB2 Rb (CST 8155S); IRAK1 Ms (SCBT sc-5288) and ASC Rb (SCBT sc-22514-R); IRAK1 Rb (CST 4504S) and NLRP3 Ms (AdipoGen AG-20B-0014-C100). At least 5000 cells were imaged for each PLA readout in iBMDM.

Antibody pairs used for PLA in BMDMs: IRAK1 Rb (Proteintech 10478-2-AP) and ASC Ms (SCBT sc-514414); IRAK1 Rb (Proteintech 10478-2-AP) and NLRP3 Ms (AdipoGen AG-20B-0014-C100); IRAK1 Ms (SCBT sc-5288) and pASC Rb (ECM Biosciences AP5631); IRAK1 Ms (SCBT sc-5288) and pJNK Rb (Invitrogen 700031); IRAK1 Ms (SCBT sc-5288) and pMKK7 Rb (CST 4171S). At least 20000 cells were imaged for each PLA readout in BMDM.

### Fluorescence Lifetime Imaging (FLIM)

Fluorescence lifetime imaging was carried out on BMDMs cultured on 8-well chamber slides (ibidi, 80826) using a Leica SP8 WLL Falcon FLIM with a 63X objective. The cells were stained with ASC and pJNK primaries (EMD Millipore 04-147 and Invitrogen 700031) and Alexa-488 and Alexa-555 secondaries, respectively. Analysis of fluorescence time decays were resolved by time-correlated single-photon counting (TCSPC) using an SPC830 acquisition board (Becker & Hickl, Berlin, Gremany). Two-photon excitation of Alexa 488 fluorophore was performed at 800 nm by a femtosecond mode-locked (80-MHz repetition rate) Mai-Tai HP pulsed multiphoton laser (Spectra Physics). Images were acquired in 1024-by 1024-pixel format, collecting in excess of 1,000 photons per pixel in 2 to 5 min, and the fluorescence transients were acquired by using SPCIMAGE software (Becker & Hickl, Berlin, Germany). The results were exported and analyzed with an image analysis protocol developed in-house using Image J imaging software.

### Antibodies for imaging

Primary – IRAK1 Rb (CST 4504S), IRAK1 Rb (Proteintech 10478-2-AP), IRAK1 Ms (SCBT sc-5288), ASC Rb (SCBT sc-22514-R), ASC Rb (CST 67824S), ASC Ms (SCBT sc-514414), ASC Ms (EMD Millipore 04-147), pASC Rb (ECM Biosciences AP5631), NLRP3 Ms (AdipoGen AG-20B-0014-C100), NLRC4 Rb (EMD Millipore 06-1125), pJNK1/2 Rb (Invitrogen 700031), IRAK4 Rb (CST 4363S), IRAK3 Rb (Thermo Scientific PA5-19969), IRAK2 Rb (Abcam ab66017), IRAK2 Ms (SCBT sc-515885), MyD88 Rb (CST 4283S), MyD88 Gt (R&D Systems AF3109), Rb IgG Isotype (CST 3900S), Ms IgG Isotype (CST 5415S), TRAF6 Rb (Bioss bs-1184R), TRAF6 Rb (Abcam ab33915), pTBK1 Rb (CST 5483S), pJNK Rb (Invitrogen 700031), pMKK4 Rb (CST 4514P), pMKK7 Rb (CST 4171S), p44/42 Rb (CST 4695P), p65 Rb (Abcam ab16502), pATF2 Ms (SCBT sc-8398), pATF2 Rb (CST 5112S), Pellino-1 (CST 31474S), TICAM2 Ms (SCBT sc-376076), pERK5 Rb (CST 3371S), pTAK1 Rb (CST 4531S), pTAB2 Rb (CST 8155S), βTrCP Rb (CST 11984S), Proteasome 20S alpha 5 Rb (Novus Biologicals NBP1-86838).

Secondary - Anti-Rabbit IgG, Alexa Fluor® 488 conjugate, (Goat, 1:1,000; ThermoFisher, catalog no. A-11034); Anti-Mouse IgG, Alexa Fluor® 488 conjugate, (Donkey, 1:1,000; ThermoFisher, catalog no. A-21202); Anti-Rabbit IgG, Alexa Fluor® 555 conjugate, (Donkey, 1:1,000; ThermoFisher, catalog no. A-31572); Anti-Rabbit IgG, Alexa Fluor® 647 conjugate, (Chicken, 1:1,000; ThermoFisher, catalog no. A-21443); Anti-Mouse IgG, Alexa Fluor® 647 conjugate, (Goat, 1:1,000; ThermoFisher, catalog no. A-21235); Anti-Mouse IgG, Alexa Fluor® 647 conjugate, (Chicken, 1:1,000; ThermoFisher, catalog no. A-21463);

### Inhibitors

Monensin (BD GolgiStop 554724), Dynasore (Cayman Chemical 14062), Thymoquinone (Sigma-Aldrich 274666), ST2825 (ApexBio A3840), JNK Inhibitor VIII (Cayman Chemical 15946), U0126 (MEK1/2 inhibitor to inhibit ERK1, Cayman Chemical 70970), XMD8-92 (ER5i, ApexBio A3943)

### Fluidigm Quantitative PCR

Quantitative PCR was carried out according to the manufacturer’s instructions using the BioMark HD system (Fluidigm), with DELTAgene primer sets designed by the manufacturer. Ct values were automatically calculated and exported from the BioMark HD system, then normalized to housekeeping gene *Hprt*.

### Cytokine ELISA

BMDMs seeded in 96-well plates were treated with the appropriate TLR ligands and for inflammasome readouts cells were triggered with ATP (5 mM). The cell culture medium was collected, and the concentrations of cytokines were determined by ELISA. Mouse IL1α, IL1ß and TNF ELISA kits were purchased from R&D Systems.

### Cytometric Bead Array (CBA)

Cytometric bead array (CBA) was performed using the mouse soluble protein master buffer kit combined with the appropriate CBA flex sets (BD Biosciences), per manufacturer’s instructions. Cell culture supernatants, serum and BAL samples were diluted 2x in assay diluent before mixing with capture beads. Flow data were collected on a BD Fortessa and analyzed with FlowJo. Data represent the median fluorescence intensity of PE on beads collected for analysis, extrapolated to protein concentration using a standard curve.

### Data Visualization

Heatmaps were generated using the matrix visualization and analysis software, Morpheus (https://software.broadinstitute.org/morpheus/). The color palettes were selected using ColorBrewer.org Cytofluorograms were generated by an ImageJ plugin JACoP.

## Supplemental Information

### Supplementary Table 1

mRNA levels assayed by Fluidigm microfluidic qPCR

### Movies

(1) iBMDM mIRAK1 mCherry cells treated with Salmonella Typhimurium expressing GFP at MOI 10

(2) iBMDM mIRAK1 mCherry cells treated with Salmonella Typhimurium expressing GFP at MOI 100

(3, 4) iBMDM mIRAK1 mCherry cells treated with 50 nM KLA and 50 nM P3C.

(5) iBMDM mIRAK1 mCherry cells treated with 100 nM KLA.

(6) iBMDM mIRAK1 mCherry cells treated with 100 nM P3C.

(7) iBMDM mIRAK1 mCherry ASC GFP cells pretreated with DMSO for 30 minutes, treated with 50 nM KLA and 50 nM P3C followed by Nigericin (10 *μ*M) for 2 hrs. Imaging starts from 5 minutes after Nigericin (10 *μ*M) treatment.

(8) iBMDM mIRAK1 mCherry ASC GFP cells pretreated with thymoquinone (25*μ*M) for 30 minutes, treated with 50 nM KLA and 50 nM P3C followed by Nigericin (10 *μ*M) for 2 hrs. Imaging starts from 5 minutes after Nigericin (10 *μ*M) treatment.

(9) iBMDM *Irak1^−/−^* ASC GFP cells pretreated with DMSO for 30 minutes, treated with 50 nM KLA and 50 nM P3C followed by Nigericin (10 *μ*M) for 2 hrs. Imaging starts from 5 minutes after Nigericin (10 *μ*M) treatment.

---

**Figure S1.**
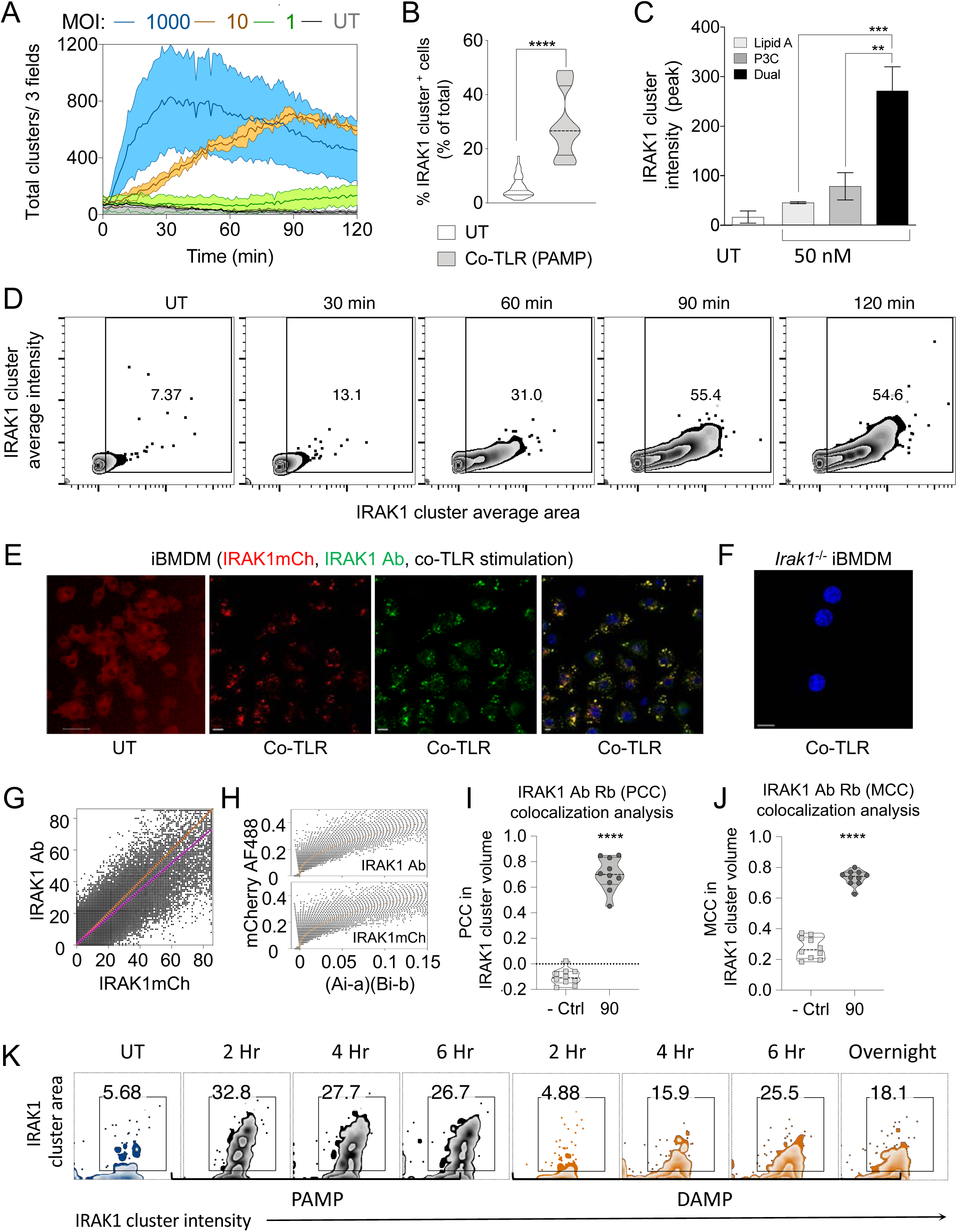
IRAK1 forms discrete intracellular clusters in response to multi-PRR activation. (A) Time course of IRAK1 clustering on multi-PRR stimulation with *Salmonella typhimurium* with the color bands showing the range. Three fields were imaged. (B-J) Kdo-2 Lipid A and P3C co-treatment (50 nM) in iBMDMs. (B) Percentage of iBMDMs showing IRAK1 clustering at 90 min, (C) IRAK1 clustering intensity in iBMDMs on single vs co-TLR stimulation at 90 min, (D) Time course of IRAK1 clustering. (E) Images of IRAK1mCherry and IRAK1 antibody staining in iBMDMs. Scale Bar: 10*μ*. (F) IRAK1 antibody (SCBT sc-5288) staining in *Irak1*^−/−^ iBMDMs. Scale Bar: 10*μ*. (G) Cytofluorogram for IRAK1 antibody (SCBT sc-5288) and IRAK1mCherry. (H) Independent Component Analysis for signals from IRAK1 antibody (SCBT sc-5288) and IRAK1 mCherry. (I) Quantification of the Pearson’s correlation coefficient (PCC) for the correlation of IRAK1 antibody (CST 4504S) staining in IRAK1 mCherry clusters in iBMDMs. (J) Quantification of the Mander’s colocalization coefficient (MCC) for the co-occurrence of IRAK1 antibody (CST 4504S) staining in IRAK1 mCherry clusters in iBMDMs. (K) Time course of IRAK1 clustering on co-stimulation of TLR4 and TLR2 with soluble biglycan (1*μ*g/mL), a DAMP or co-TLR stimulation with the PAMPs - P3C and Kdo-2 Lipid A (50 nM each). Data are represented as median. (Panels B, C) Kolmogorov-Smirnov test. (Panels I, J) Each data point represents value from a randomly selected non-over lapping field of cells. Paired t test. p = 0.1234 (ns), 0.0332(*); 0.0021 (**); 0.0002 (***); < 0.0001 (****). Data shown are representative of at least two independent experiments.

**Figure S2.**
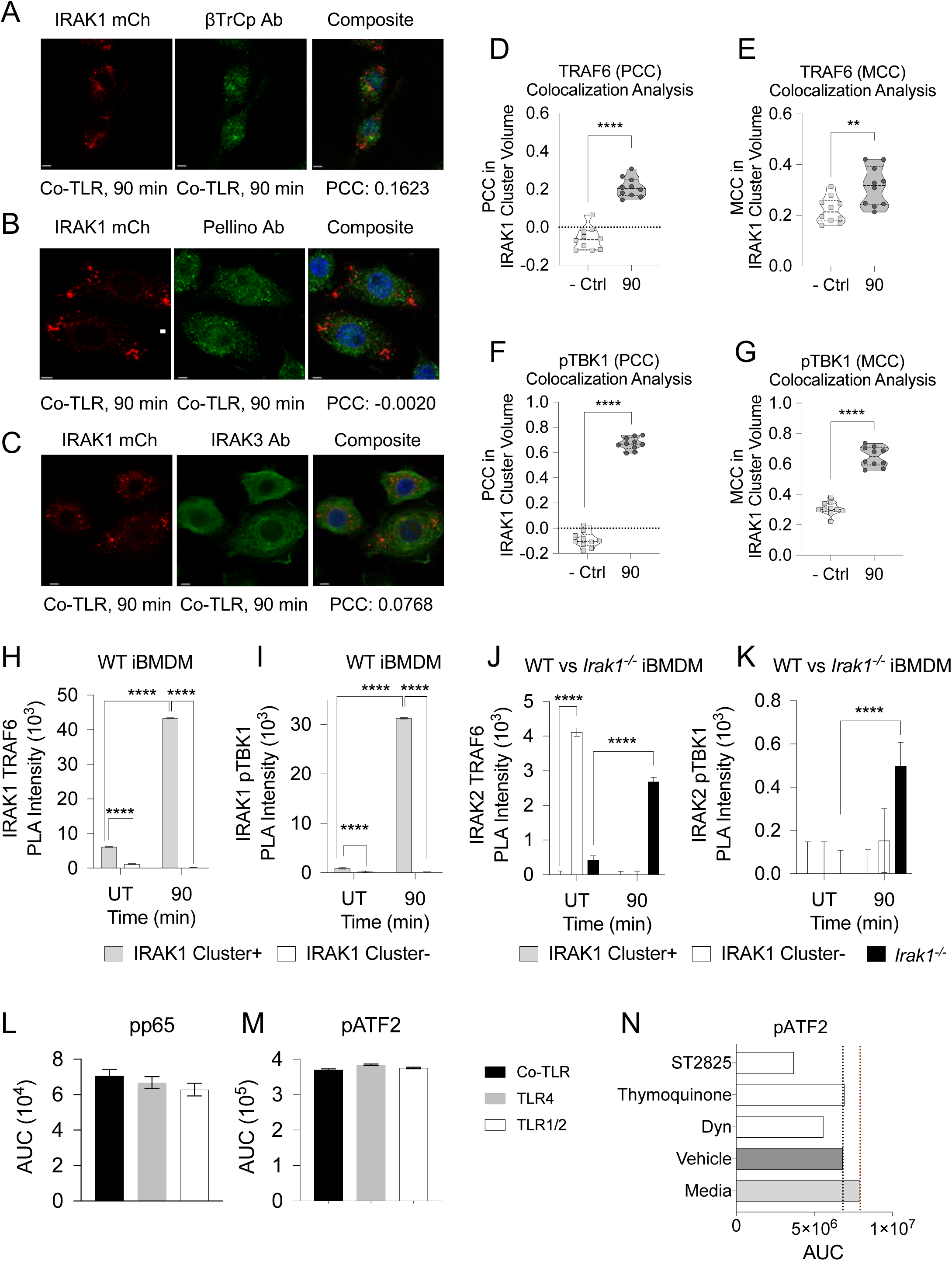
Characterizing IRAK1 clusters. (A-K) Kdo-2 Lipid A and P3C co-treatment (50 nM each) in iBMDMs. Co-staining of IRAK1 clusters and (A) ßTrCp, (B) pellino, and (C) IRAK3. Scale Bar: 10*μ*. (D) Quantification of the Pearson’s correlation coefficient (PCC) for the correlation of TRAF6 antibody staining in IRAK1 clusters. (E) Quantification of the Mander’s colocalization coefficient (MCC) for the co-occurrence of TRAF6 antibody staining in IRAK1 clusters in iBMDMs. (F) PCC and (G) MCC for pTBK1 antibody staining in IRAK1 clusters. (H-K) Proximity ligation assay in iBMDMs. (H) IRAK1-TRAF6 PLA, (I) IRAK1-pTBK1 PLA, (J) IRAK2-TRAF6 PLA, (K) IRAK1-pTBK1 PLA. (L, M) Single or co-TLR stimulation of TLR4 and TLR1/2 with Kdo-2 Lipid A and P3C (50 nM each). Quantification of (L) pp65 nuclear translocation (M) pATF2 nuclear translocation. (N) Quantification of pATF2 nuclear translocation on co-TLR stimulation of TLR4 and TLR1/2 with Kdo-2 Lipid A and P3C ± 30 min pre-treatment with dynasore, an internalization inhibitor, 20*μ*M, ST2825, a MyD88 dimerization inhibitor, 20*μ*M, and thymoquinone, an IRAK1 kinase inhibitor, 25*μ*M. (D-G) Data are represented as mean ± SD. (H-N) Data are represented as median ± MAD. (Panels D-G) Each data point represents value from a randomly selected non-over lapping field of cells. Paired t test. (Panels H-M) Tukey’s multiple comparisons test. (Panel N) Dunnett’s multiple comparisons test. p = 0.1234 (ns), 0.0332(*); 0.0021 (**); 0.0002 (***); < 0.0001 (****). Data shown are representative of at least two independent experiments.

**Figure S3.**
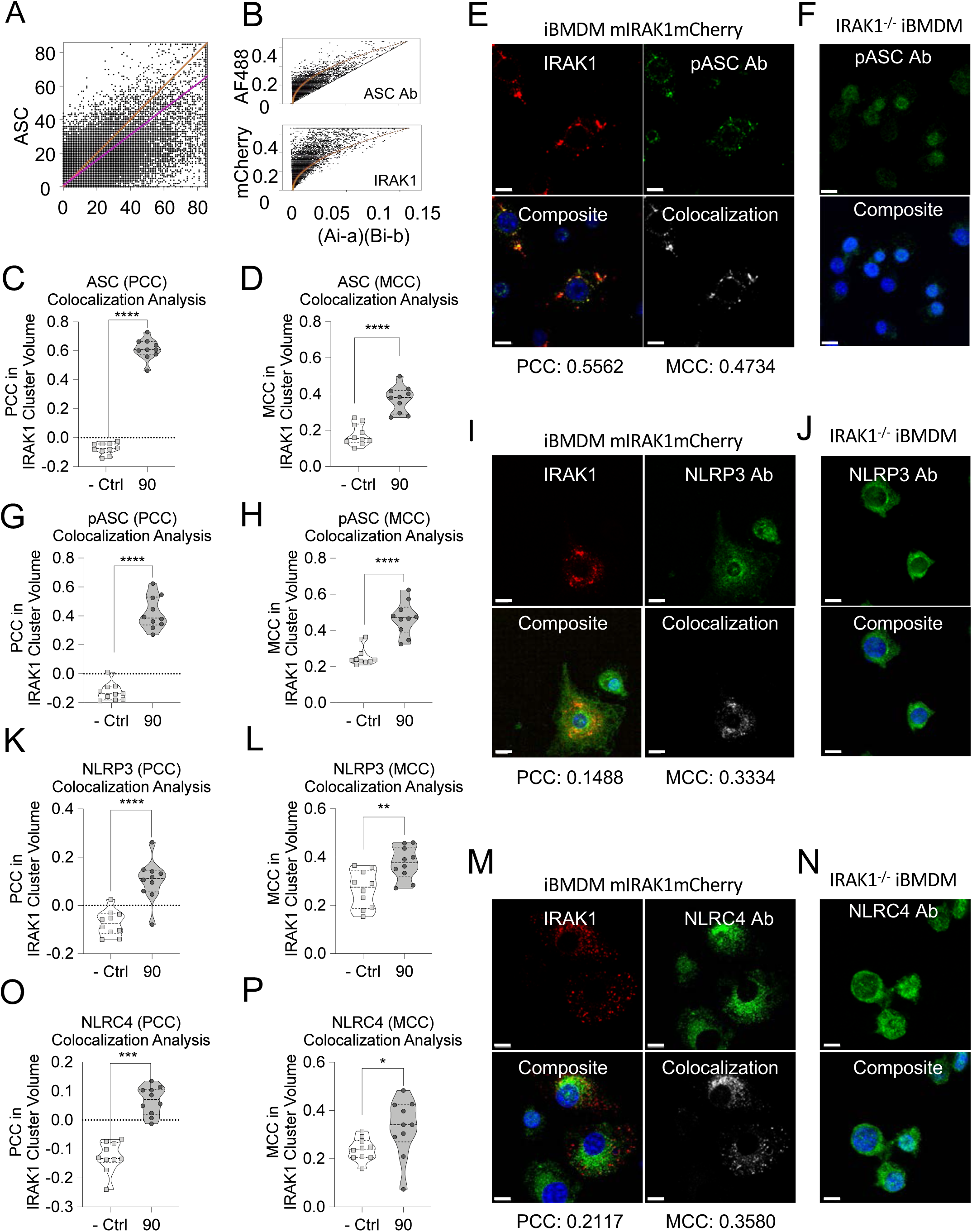
IRAK1 clusters recruit inflammasome components in iBMDMs. Kdo-2 Lipid A and P3C co-treatment (50 nM each) in iBMDMs. (A) Cytofluorogram for ASC antibody (SCBT sc-22514-R) and IRAK1mCherry. (B) Independent Component Analysis for signals from ASC antibody (SCBT sc-22514-R) and IRAK1 mCherry. (C) Quantification of the Pearson’s correlation coefficient (PCC) for the correlation of ASC antibody (SCBT sc-22514-R) staining in IRAK1 clusters. (D) Quantification of the Mander’s colocalization coefficient (MCC) for the co-occurrence of ASC antibody (SCBT sc-22514-R) staining in IRAK1 clusters in iBMDMs. (E) Co-staining of IRAK1 clusters and pASC. (F) pASC staining in *Irak1*^−/−^ iBMDMs. (G) PCC and (H) MCC for pASC antibody staining in IRAK1 clusters. (I) Co-staining of IRAK1 clusters and NLRP3. (J) NLRP3 staining in *Irak1*^−/−^ iBMDMs. (K) PCC and (L) MCC for NLRP3 antibody staining in IRAK1 clusters. (M) Co-staining of IRAK1 clusters and NLRC4. (N) NLRC4 staining in *Irak1*^−/−^ iBMDMs. (O) PCC and (P) MCC for NLRC4 antibody staining in IRAK1 clusters. (Panels C, D, G, H, K, L, O, P) Each data point represents value from a randomly selected non-over lapping field of cells. Paired t test. Automated imaging of at least 2 (Panels F, J, N) fields of cells. Data are represented as mean ± SD. p = 0.1234 (ns), 0.0332(*); 0.0021 (**); 0.0002 (***); < 0.0001 (****). Data shown are representative of at least two independent experiments.

**Figure S4.**
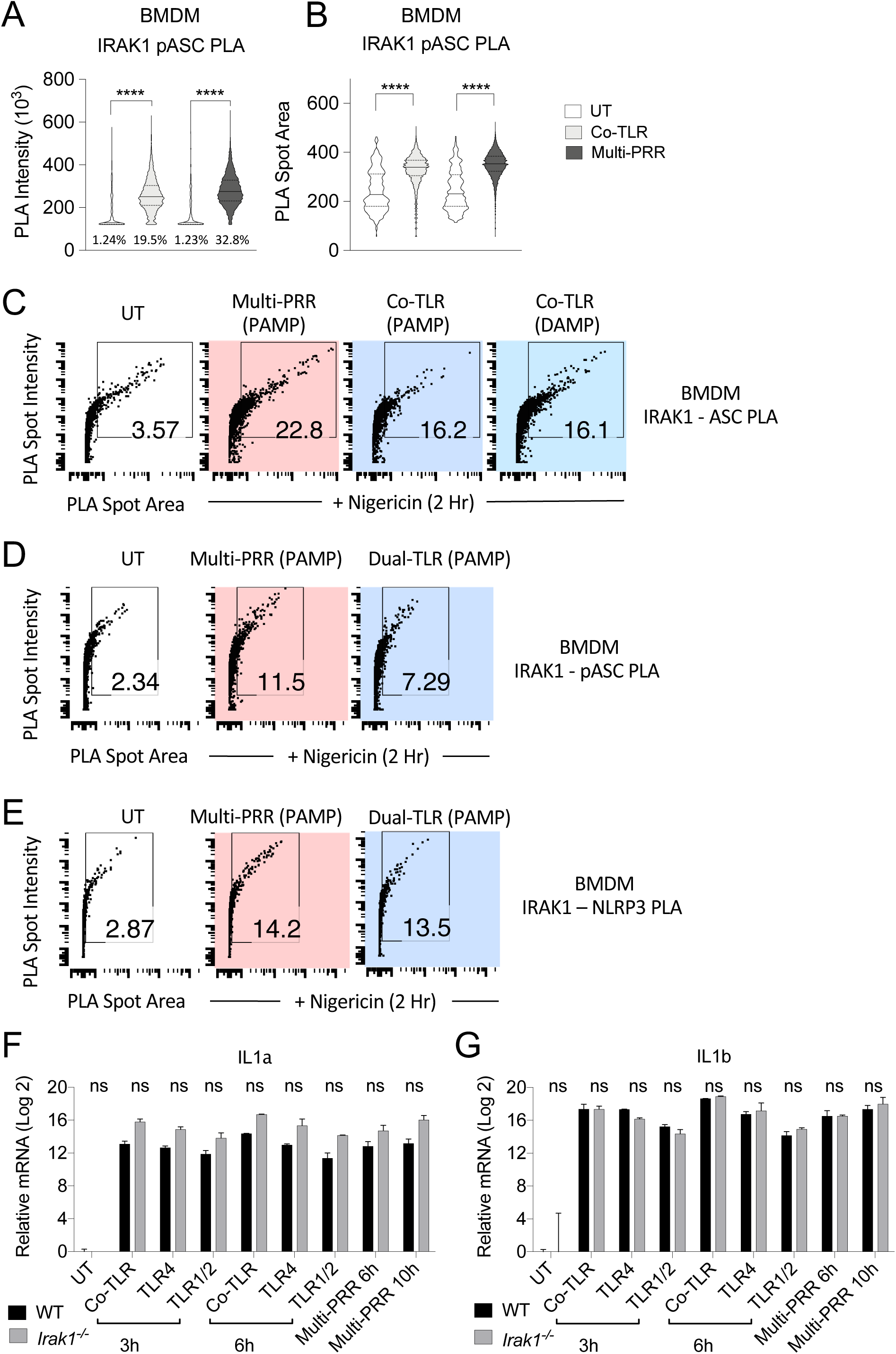
IRAK1 clusters recruit inflammasome components in BMDMs. Proximity ligation assay in BMDMs. At least 20,000 cells imaged for all graphs. Co-TLR stimulation with Kdo-2 Lipid A and P3C co-treatment (500 nM). Multi-PRR stimulation using heat-killed *Yersinia pseudotuberculosis* (MOI 100). (A) IRAK1-pASC PLA intensity, (B) IRAK1-pASC PLA spot area. (C-E) Co-TLR (PAMP: Kdo-2 Lipid A and P3C, DAMP: soluble biglycan) or multi-PRR priming (heat-killed *Yersinia pseudotuberculosis*, MOI 100) followed by Nigericin trigger (10 *μ*M) for 2h (C) IRAK1-ASC PLA, (D) IRAK1-pASC PLA, (E) IRAK1-NLRP3 PLA. qPCR quantification of (F) *IL1a* and (G) *IL1b* in BMDMs primed with co-TLR stimulation using 500 nM of both Kdo-2 Lipid A and P3C or single-TLR stimulation with either 1*μ*M Kdo-2 Lipid A or 1*μ*M P3C or with heat-killed Mean ± SD from at least 4 wells. (Panels A-B) Kolmogorov-Smirnov test. (Panels F, G) Šídák’s multiple comparisons test. p = 0.1234 (ns), 0.0332(*); 0.0021 (**); 0.0002 (***); < 0.0001 (****). Data shown are representative of at least two independent experiments.

**Figure S5.**
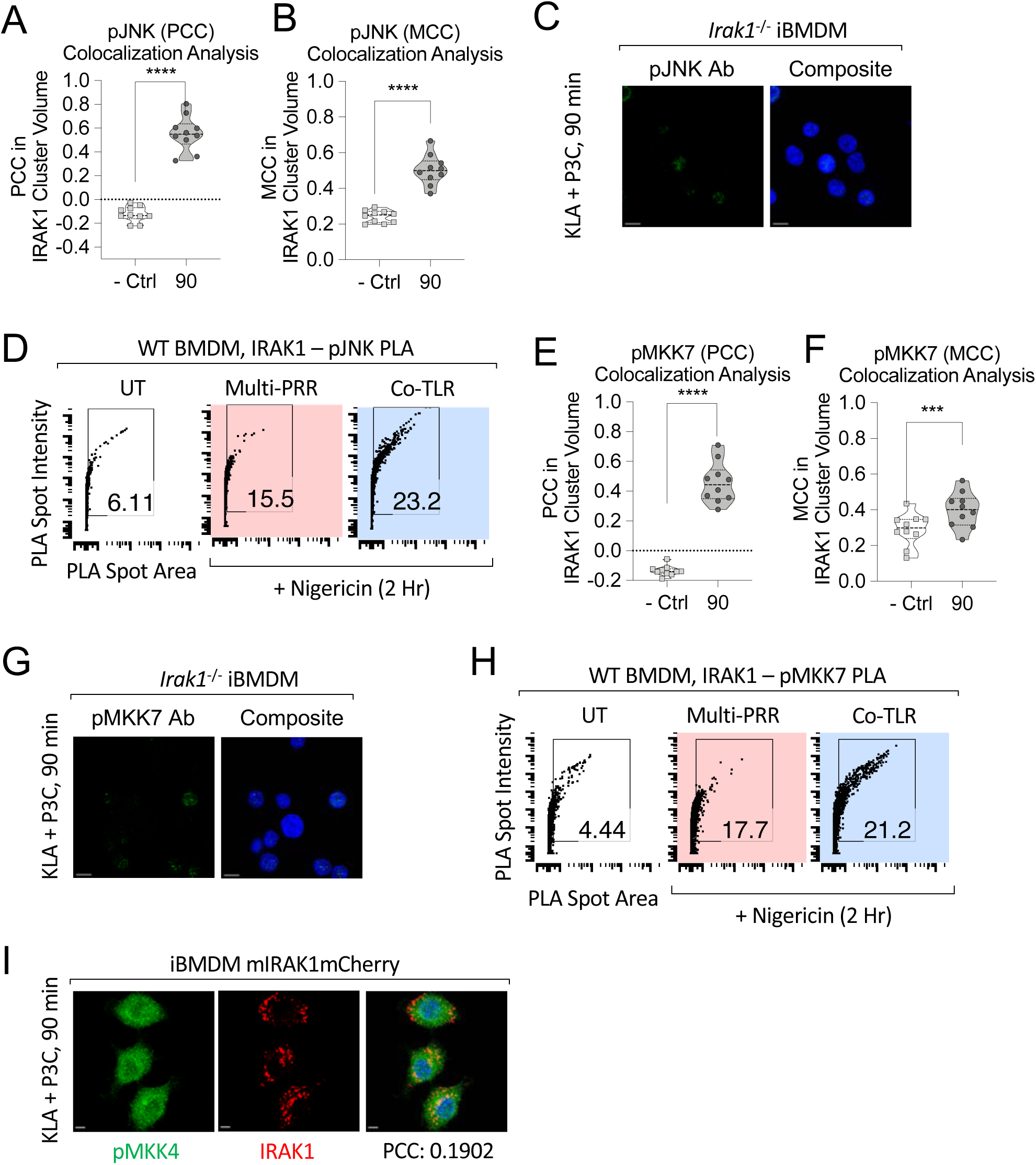
IRAK1 clusters are dependent on a recruited JNK MAPK cascade. (A-C, E-G, I) Kdo-2 Lipid A and P3C co-treatment (50 nM each) in iBMDMs. (D, H) Co-TLR (PAMP: Kdo-2 Lipid A and P3C, 50 nM each) or multi-PRR priming (heat-killed *Yersinia pseudotuberculosis*, MOI 100) followed by Nigericin (10 *μ*M) trigger for 2h. (A) Quantification of the Pearson’s correlation coefficient (PCC) for the correlation of pJNK antibody staining in IRAK1 clusters. (B) Quantification of the Mander’s colocalization coefficient (MCC) for the co-occurrence of pJNK antibody staining in IRAK1 clusters. (C) pJNK staining in *Irak1*^−/−^ iBMDMs. (D) IRAK1-pJNK PLA. (E) PCC and (F) MCC for pMKK7 antibody staining in IRAK1 clusters. (G) pMKK7 staining in *Irak1*^−/−^ iBMDMs. (H) IRAK1-pMKK7 PLA in BMDMs. (I) pMKK4 staining in iBMDMs mIRAK1mCherry. (Panels A, B, E, F) Each data point represents value from a randomly selected non-over lapping field of cells. Paired t test. Data are represented as mean ± SD. Automated imaging of at least 2 (Panels C, G, I) fields of cells. At least 20000 (Panels D, H) cells imaged. p = 0.1234 (ns), 0.0332(*); 0.0021 (**); 0.0002 (***); < 0.0001 (****). Data shown are representative of at least two independent experiments.

